# Direct encoding of new visual concepts in early visual cortex

**DOI:** 10.1101/2025.10.07.680862

**Authors:** Svenja Klinkowski, Anna Seewald, Björn Fath, Vanessa Kasties, Panagiotis Iliopoulos, Silke Schmidt, Franziska Voss, Michael Erb, Klaus Scheffler, Steffen Gais, Svenja Brodt

## Abstract

Neocortical memories are assumed to rely on slow systems consolidation through hippocampal offline reactivation. However, recent studies show rapid neocortical integration of new memories for information associated with prior knowledge. Testing the limits of rapid neocortical learning, we tracked memory formation across 24h with functional and diffusion-weighted MRI in subjects acquiring either conceptual or detailed knowledge of novel objects from identical visual stimulation. Concept learning-induced rapid, 24h-stable changes in strength and pattern of functional early visual cortex responses. These were co-localized with microstructural changes, correlated positively with categorization performance and negatively with context recognition. Detailed item-context learning elicited analogous changes in posterior parietal areas. Results show that the neocortex can rapidly learn without prior knowledge and that concept learning can occur in early sensory regions that process corresponding visual features. Thus, we substantiate the possibility of direct or parallel neocortical encoding without offline abstraction from individual episodes.

## Introduction

The architecture of the neocortex is tailored to store large amounts of information (Brunel, 2016). Particularly for declarative memory, the neocortex has traditionally been considered a slow learning system, which requires time and a large number of repetitions to establish a memory trace. These repetitions are assumed to originate from spontaneous hippocampal replay of episodic memories during offline states such as sleep (Brodt et al., 2023; Frankland and Bontempi, 2005). Based on this idea, a central proposition is that neocortical memories become abstracted from episodic detail in the process (Kumaran and McClelland, 2012; McClelland et al., 1995; Moscovitch et al., 2016; Winocur and Moscovitch, 2011) with semanticized representations relying on higher-order associative neocortical areas such as medial posterior parietal cortex (mPPC) and medial prefrontal cortex (Binder and Desai, 2011; Tompary and Davachi, 2017; Tse et al., 2011). However, this framework of complementary memory systems and systems memory consolidation has recently been challenged in several important ways (Gilboa and Moscovitch, 2021; Renoult et al., 2019).

Most significantly, neocortical memory traces can be established much more rapidly than previously assumed. Animal studies have shown that optogenetic reactivation of neocortical engram cells can establish hippocampal independence within 24 h (Cowansage et al., 2014; de Sousa et al., 2019). In humans, evidence for rapid neocortical memory formation comes from studies employing repeated learning of novel associations, e.g. between words, colors, objects, or locations (Brodt et al., 2018; Brodt et al., 2016; Flanagin et al., 2023; Himmer et al., 2019). In those studies, learning induced lasting changes in neocortical memory networks within a single session, similar to what has previously been expected to occur only over days or weeks during systems consolidation (McClelland et al., 1995). We previously showed the rapid development of a neocortical engram during learning in the mPPC using diffusion-weighted MRI (dMRI), which assesses microstructural plasticity (Brodt et al., 2018). However, it is currently unknown which types of memories can be acquired rapidly by the neocortex. Memory tasks used in previous studies employed novel associations between known items (e.g., associating words or images of known objects or people) (Sommer, 2017; van Kesteren et al., 2013; van Kesteren et al., 2010), integration of new places into spatial schemas (Tse et al., 2007; Tse et al., 2011), or fast mapping (Coutanche and Thompson-Schill, 2014; Merhav et al., 2015; Sharon et al., 2011). Besides reactivation via repetition and repeated retrieval (Antony et al., 2017; Brodt and Gais, 2020), prior knowledge has been proposed to facilitate the rapid integration of new elements within existing neocortical memory networks (Hebscher et al., 2019b; Sekeres et al., 2024; Tse et al., 2011). Testing whether novel information that is entirely unrelated to prior knowledge can also be learned rapidly by the neocortex is the first aim of the present study.

Another challenge to the current memory systems framework is recent evidence indicating that instead of a sequential systems consolidation process, memory encoding of detailed and generalized representations can occur in parallel in different memory systems, which for both can entail neocortical regions (Cowansage et al., 2014; Gilboa and Moscovitch, 2021). This would in particular affect our understanding of semantic memory, as traditionally it is assumed to form gradually in the neocortex through repeated offline reactivation of hippocampal episodic representations (Gilboa and Moscovitch, 2021; Krenz et al., 2023; Navarro Lobato et al., 2023). With parallel neocortical encoding, concept formation could directly occur within the neocortex, putatively including modality-specific sensory as well as higher-order integration areas (Binder and Desai, 2011; Bracci and Op de Beeck, 2016; Goltstein et al., 2021; Popham et al., 2021; Yee and Thompson-Schill, 2016). Of note, we refer to the definition by Gilboa and Marlatte (Gilboa and Marlatte, 2017) of a concept as a “mental representation denoting a class of items or entities that belong to a particular category”. Here, we specifically tested whether detailed and conceptual memories can be encoded directly into the neocortex. We hypothesized that both memory types differ in their temporal and spatial recruitment of memory networks.

Thus, in the present study we examined whether rapid neocortical learning might be a much more common phenomenon than previously assumed, occurring for novel concepts as well as for novel detailed item-context memories. To investigate whether such memories can be rapidly established in the neocortex in the absence of prior knowledge, we followed the evolution of memories for entirely novel, abstract visual stimuli in human subjects across 24 h with the help of multimodal MRI (Fig. 1A). Two groups of participants (N_total_=80) were presented with identical visual stimuli in form of blob-shaped, colored “aliens” (items) in front of different planets (contexts). One group was instructed to remember individual item-context combinations (detail group), while the other (concept group) was instructed to remember and identify families of aliens (categories) with similar shape and color (Fig. 1B).

**Fig. 1.**
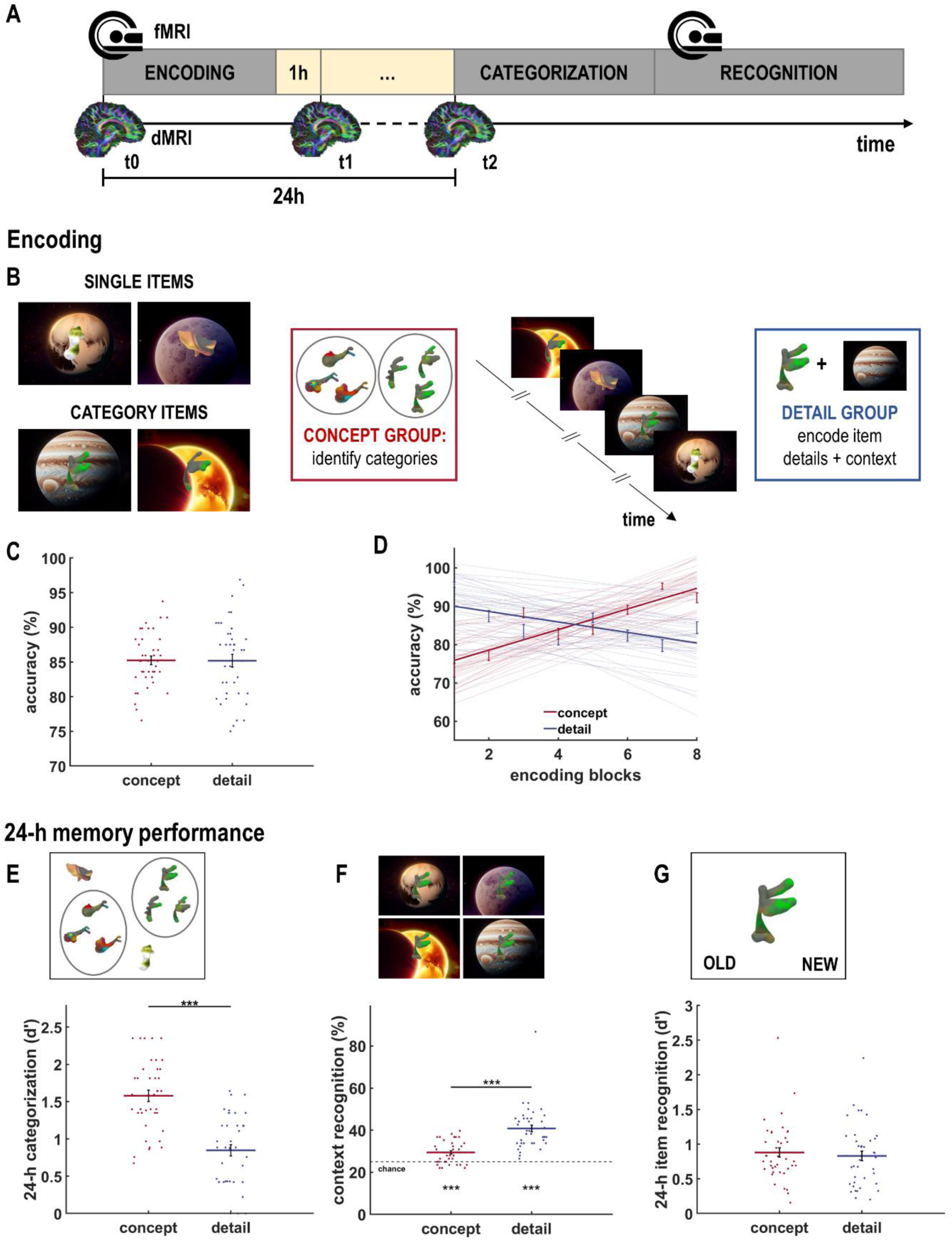
Experimental Design and Behavioral Results. (**A**) On the first day, participants performed the encoding task during fMRI. 24 h later, categorization performance was assessed, and item-context recognition memory was tested during fMRI. dMRI was conducted at three time points: before encoding as baseline (t0), 1 h after the end of encoding (t1) and before the memory tests on the second day (t2). (**B**) During encoding, two types of abstract visual stimuli were presented: single items with unique features and category items sharing a similar shape and colour scheme. Items (“aliens”) were presented with a planet as background (context). Two groups of participants (N=2×40) were assigned different instructions on how to encode the same visual stimuli. The concept group was told to find similarities between items and to categorize them into “families”, while the detail group was told to encode specific item details in combination with the context. (**C**) The concept and detail groups did not differ in their overall performance throughout the encoding task, indicating similar task difficulty. (**D**) The encoding task consisted of 8 encoding blocks, each with 16 trials. Throughout the task, accuracy in the detail group decreased, corresponding to the increasing number of intervening item-context associations, whereas the concept group improved in detecting categories. (**E**) During the categorization task, participants were asked to judge whether novel items belonged to one of the learned categories. While both groups performed above chance level, the concept group performed significantly better at categorizing new items (d’_concept_=1.58, d’_detail_=0.85, *t*_78_=6.91, *p*<0.001). (**F** and **G**) The item-context recognition task consisted of two phases: (F) During context recognition, the participants had to identify in which context items were presented during encoding with a four-alternative forced choice. While both groups performed above chance level (concept: *t*_39_=5.54, *p*<0.001; detail: *t*_39_=9.80, *p*<0.001), the detail group remembered significantly more item-context combinations (correct recognition: cr_concept_=29.5%, cr_detail_=40.8%, *t*_78_=-6.31, *p*<0.001). (G) During item recognition, participants judged whether items presented without context were old or new. Recognition performance did not differ between the two groups (d’_concept_=0.88, d’_detail_=0.83, *t*_78_=0.51, *p*=0.61), indicating similar memory of individual items. Red, concept group; blue, detail group; bars, mean; error bars, SEM; dots, individual data points; *, *p*<0.05; ***, *p*<0.001.

To assess learning-induced brain plasticity, we use a combination of functional and diffusion-weighted MRI previously shown to detect rapid neocortical engram formation (Brodt et al., 2018). Microstructural plasticity is measured via changes in mean diffusivity (MD) which reflect changes in the degree of hindrance or restriction of water diffusion within a voxel. The origin of these changes were explored with Neurite Orientation Dispersion and Density Imaging (NODDI), a biophysical diffusion model disentangling the relative contributions of CSF, neurites and cell somata (Zhang et al., 2012).

## Results

### Different encoding goals induce conceptual vs. detailed memory of the same stimuli

Overall performance during the encoding task did not differ between the two groups despite the different encoding instructions (mean accuracy: 85%; *t*_78_=0.05, *p*=0.96; Fig. 1C), nor did the subjective perception of task difficulty (*t*_78_=0.49, *p*=0.63). However, over the course of the task, accuracy in the concept group increased, corresponding to the accumulation of category evidence (mean slope=2.69, *t*_39_=14.09, *p*<0.001), while performance in the detail group decreased (mean slope=-1.38, *t*_39_=-7.46, *p*<0.001) corresponding to the increased number of item-context associations (difference between slopes: *t*_78_=15.33, *p*<0.001; Fig. 1D). 24 h after encoding, the concept group performed significantly better in assigning novel, unseen stimuli to the previously encoded categories (Fig. 1E), whereas the detail group was better at remembering the specific item-context combinations (Fig. 1F). Hence, the groups preferentially encoded different aspects of the same stimuli, leading to differences in memory quality after 24 h.

### Rapid emergence of category-sensitive functional plasticity in early visual cortex

To investigate how memories for entirely novel concepts develop, we compared changes in brain activity over category repetitions between the concept group and the detail group during encoding, thus disentangling category learning from general memory effects, visual stimulation, and time. With increasing category evidence, activity in secondary visual cortex (V2) – ventrally extending along the lingual gyrus and dorsally along the cuneus – linearly increased in the concept but not in the detail group (increase × concept-detail, *p*_FWE_<0.05; Fig. 2A, B and table S1). Moreover, only the concept group displayed higher activity within this occipital cluster for repeated category items compared with items that did not belong to a category (group × stimulus type, *F*_1,78_=12.18, *p*<0.001, Fig. 2C). This category-specific activity was positively correlated with improved categorization performance 24 h later specifically in the concept group (*r_concept_*=0.39, *p*=0.01; *r_detail_*=0.02, *p*=0.91; comparison between groups: *z*_btw_=1.68, *p*=0.046; Fig. 2D). These results show that category learning induces the rapid emergence of a category-specific early visual cortex response that predicts successful category memory 24 h later.

**Fig. 2.**
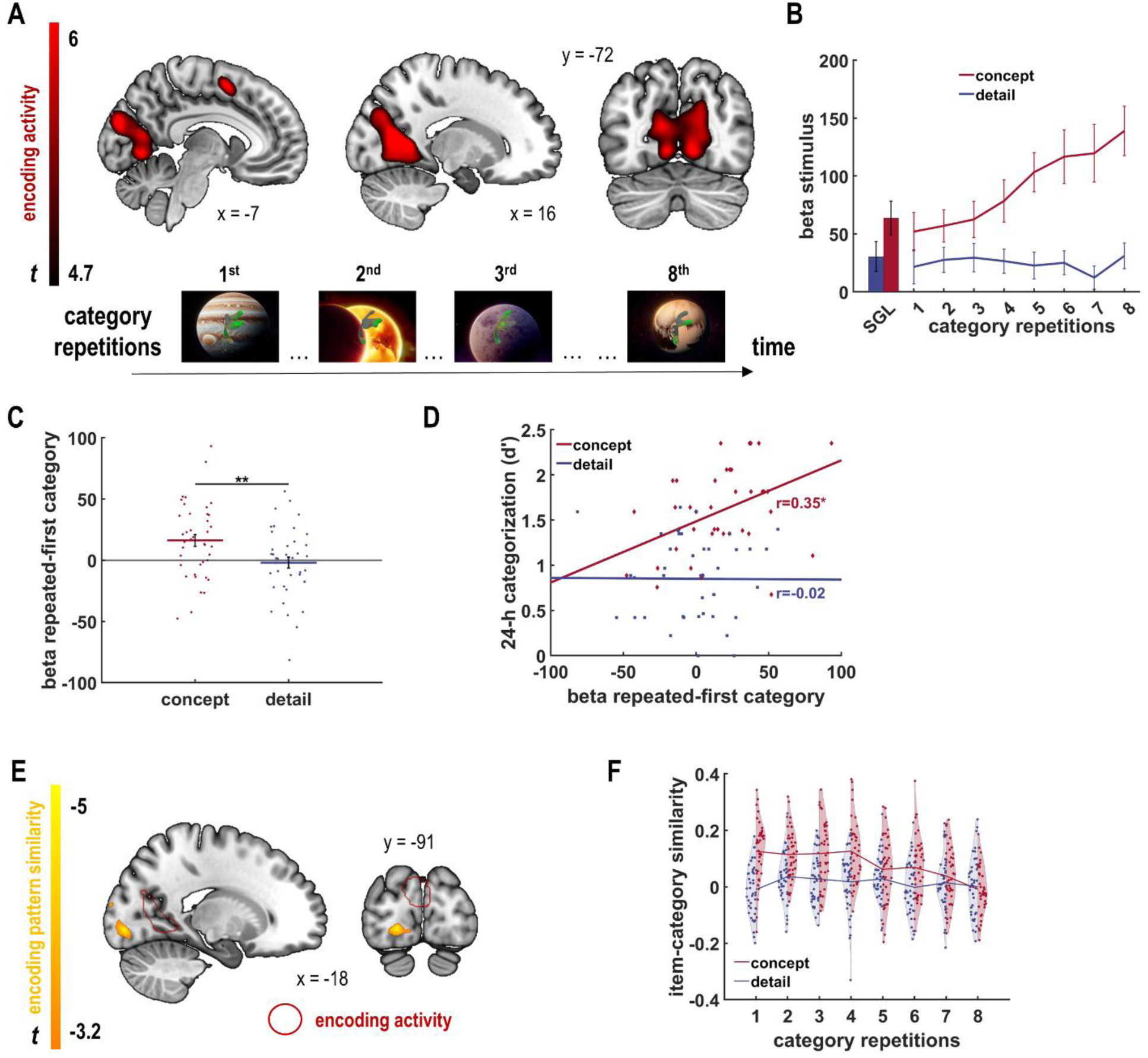
Rapid formation of visual concepts in early visual cortex during encoding. (**A** and **B**) During the encoding task, the concept group showed an increase in functional activity in early visual cortex, in particular in V2, with increasing category evidence, compared with the detail group (increase over category repetitions × concept>detail, full-volume family-wise error (FWE) corrected *p*_FWE_<0.05). (B) This group-specific increase in activity was confined to category stimuli and there was no significant group difference in activity for “single” items that were presented only once (*t*_78_=1.70, *p*=0.09; “SGL”). (**C**) In the activity cluster from (A), the concept group displayed higher category-specific encoding activity than the detail group when comparing repeated category items to the first occurrence of a category including single items. (**D**) This category-specific activity predicted improved categorization performance 24 h later, but only in the concept group. (**E** and **F**) In an additional whole-brain analysis, we searched for regions where the activity pattern representing individual stimuli changed with regard to the average pattern representing a category. During encoding, pattern similarity between individual category members and the average category pattern decreased in the concept group over category repetitions at the occipital pole (decrease in item-category similarity × concept>detail, region depicted at *p*_uncorr_<0.001, peak activity full-volume corrected at *p*_FWE_<0.05; purple-yellow). No such decrease in item-concept-similarity was found for the detail group, which focused on differences between items. There was no significant group-specific increase in item-concept-similarity over concept repetitions in any region. A control analysis comparing each category item to the mean pattern of the other categories yielded no significant clusters. Red cluster, concept-specific increase during encoding from (A); red, concept group; blue, detail group; dots, individual data points; *, *p*<0.05; **, *p*<0.01.

Complementing these univariate analyses, we investigated the neural representation of newly developing concepts by assessing pattern similarity. Specifically, we correlated the neural activity pattern evoked by each category item with the average evoked pattern of all items of the same category. We could thus follow concept development over the eight item presentations of each category. A whole-brain mixed model searchlight showed a significant linear decrease in item-concept-similarity over concept repetitions at the occipital pole (peak in primary visual cortex, V1) in the concept but not in the detail group (increase × concept>detail, *t*_636_=-4.83, *p_FWE_*<0.05; Fig. 2E, F and table S2). In line with ideas of predictive coding, this finding can be a result of increasing feedback modulation from a downstream area gathering category information (Gilbert and Li, 2013). Moreover, it corroborates the idea that during concept acquisition adaptive responses to complex stimuli can occur rapidly and very early along the visual processing hierarchy (Reber et al., 1998).

### Changes in early visual cortex are stable and co-occur with microstructural plasticity

Next, we assessed the stability of these early functional changes in a follow-up recognition task after 24 h. When presented with the learned stimuli, the concept group showed higher task-related activity in the occipital fusiform gyrus compared with the detail group despite identical visual stimulation and instruction in both groups (task>baseline × group, *p*_FWE_<0.05, Fig. 3A and table S3). This cluster overlapped with the cluster showing the group-specific increase in activity during encoding (region of encoding activity: *F*_1,78_=4.88, *p*=0.03; Fig. 3A, B). In this cluster, a higher category response during encoding was associated with higher activity during context recognition 24 h later specifically in the context group (*r_concept_*=0.42, *p*=0.008; *r_detail_*=-0.17, *p*=0.30; comparison between groups: *z_btw_*=2.64, *p*=0.004; Fig. 3C) showing that the early visual cortex response established during concept learning is retained over a day. This concept-related visual cortex reactivation, however, interfered with detailed context memory as it was negatively correlated with item-context recognition in the concept group (*r_concept_*=-0.34, *p*=0.03; *r_detail_*=0.03, *p*=0.85; comparison between groups: *z_btw_*=-1.65, *p*=0.049; Fig. 3D). Since the encoded stimuli are presented again during the recognition task, this activity suggests an automatic concept reactivation in the concept group, which then interferes with context memory.

**Fig. 3.**
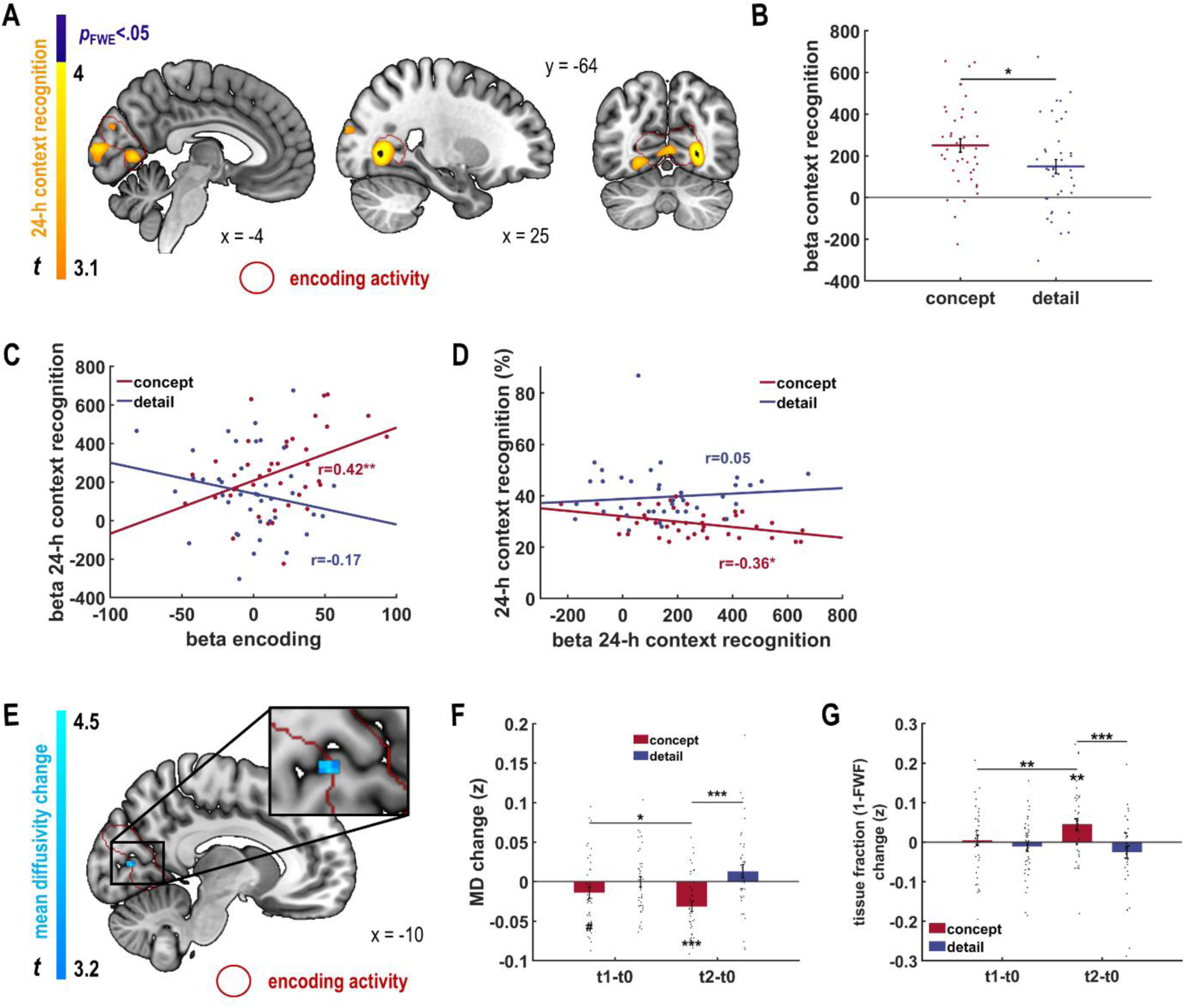
Stability of functional changes and concept-specific micro-structural changes in early visual cortex after 24 h. (**A**) 24 h after encoding, the concept group persistently displayed increased functional activity in early visual cortex compared with the detail group, during stimulus presentation in the recognition task (stimulation>baseline × concept>detail, *p*_uncorr_<0.001, orange-yellow; peak: full-volume corrected *p*_FWE_<0.05, purple). This activity overlapped with the cluster from encoding (red; see Fig. 2A). (**B** and **C**) The renewed presentation of the same stimuli 24 h after encoding elicited a reactivation of the category-specific activity in early visual cortex only in the concept group. Activity correlated with category-specific activity in the same region in the concept group, but not in the detail group (see Fig. 2A), suggesting an automatic reactivation of concept information. (**D**) Correspondingly, higher reactivation of this category-specific activity impaired item-context recognition in the concept group. (**E**) At t2, after 24 h, the concept group showed a stronger decrease in mean diffusivity (MD) in the intracalcarine cortex (t2<t0 × concept>detail, *p*_uncorr_<0.001, k>10; blue), in a cluster overlapping with the functional activity increase shown by this group during encoding (red, see Fig. 2A). (**F**) This effect can be attributed to decreased diffusivity in the concept group (concept group: t2 − t0: *t*_39_=-5.01, *p*<0.001), and no decrease in MD was observed in the detail group (detail group: t2 − t0: *t*_39_=2.65, *p*=0.01). The decrease in MD started to develop already at t1, 1 h after encoding (concept group: t1 − t0: *t*_39_=-1.96, *p*=0.057), indicating rapid and persistent microstructural changes specific to concept-learning. (**G**) The cluster showing concept-learning specific MD changes also displayed a group-specific increase in the tissue volume fraction (1-FWF) after 24 h indicating a higher relative contribution of brain tissue vs. CSF compartments after learning. Red, concept group; blue, detail group; bars, mean; error bars, SEM; dots, individual data points; *, *p*<0.05; **, *p*<0.01; ***, *p*<0.001.

Whereas fMRI can only assess learning-induced functional response changes during stimulus processing, dMRI captures changes in tissue microstructure that persist even in an offline state. These changes can not only be detected in white matter (Sampaio-Baptista et al., 2020; Scholz et al., 2009) but have been shown to indicate memory-related gray matter plasticity through decreases in averaged, non-directional molecule movement (mean diffusivity, MD) (Brodt et al., 2018; Sagi et al., 2012; Stee et al., 2023; Tavor et al., 2020). Therefore, dMRI can provide an indication of the contribution of different brain systems to memory storage (Brodt et al., 2018; Stee et al., 2023; Tavor et al., 2020). To assess the evolution of group-specific microstructural plasticity, we compared MD immediately before (t0) and 24 h after learning (t2) between both groups in order to attribute the observed changes to the specific type of memory. In a whole-brain, voxel-wise comparison, the largest cluster of MD decrease specific to the concept group was found in the left intracalcarine cortex, overlapping with the region that showed the concept-specific activity increase during encoding (t2<t0 × concept>detail, *p*_uncorr_<0.001; Fig. 3E, F and table S4A). To further explore the origin of this learning-induced increased hindrance of water diffusion, we employed NODDI, a biophysical model attributing the measured signal to three different tissue compartments, namely free water (modeling diffusion in CSF), intracellular (modeling diffusion in and around neurites) and extracellular space (modeling diffusion in and around cell somata) (Zhang et al., 2012). The cluster showing significant changes in MD additionally showed a concept-learning specific increase in the tissue volume fraction (1-FWF) after 24h (difference t2 − t0: concept – detail *t*_78_=-3.46, *p*<0.001; Fig. 3G), indicating a higher relative contribution of brain tissue vs. CSF compartments after learning. There were no significant changes in neurite density (NDI) and orientation dispersion index (ODI) (NDI: difference t2 − t0: concept – detail *t*_78_=-0.92, *p*=0.36; ODI: difference t2 − t0: concept – detail *t*_78_=-0.32, *p*=0.75), indicating that there was no systematic alteration in the relative contributions of the intracellular vs. extracellular tissue compartment or the complexity of neurite orientation. Thus, learning novel visual concepts induced microstructural plasticity in early visual cortex within 24 h, which was co-localized with the rapid learning-induced functional changes and driven by an increase in tissue density.

### Rapid acquisition of novel detailed information in mPPC

Whereas the occipital cortex showed gradual changes specific to the concept group, changes in encoding-related activity common to both groups were located further down the visual processing stream at the border between occipital and parietal lobes. Here, we observed an increase in activity already from the first to the second presentation of a category, which then remained constant with additional category repetitions in both groups (conjunction of concept & detail group: repeated>first category items, *p*_FWE_<0.05; Fig. 4A, B and table S5A), indicating a rapid change in the functional response to learned stimuli. This conjunction cluster was reactivated in the recognition task, which presented the learned stimuli 24 h later (Fig. 4C, both groups: *r*=0.31, *p*=0.001; *r_detail_*=0.30; *r_concept_*=0.31; *z*_btw_=-0.06, *p*=0.48). Reactivation was positively associated with context recognition performance in the detail group (*r_detail_*=0.37, *p*=0.02; *r_concept_*=-0.16, *p*=0.31; comparison between groups: *z*_btw_=-2.38, *p*=0.009; Fig. 4D), which indicates a particular role of the mPPC for episodic memory.

**Fig. 4.**
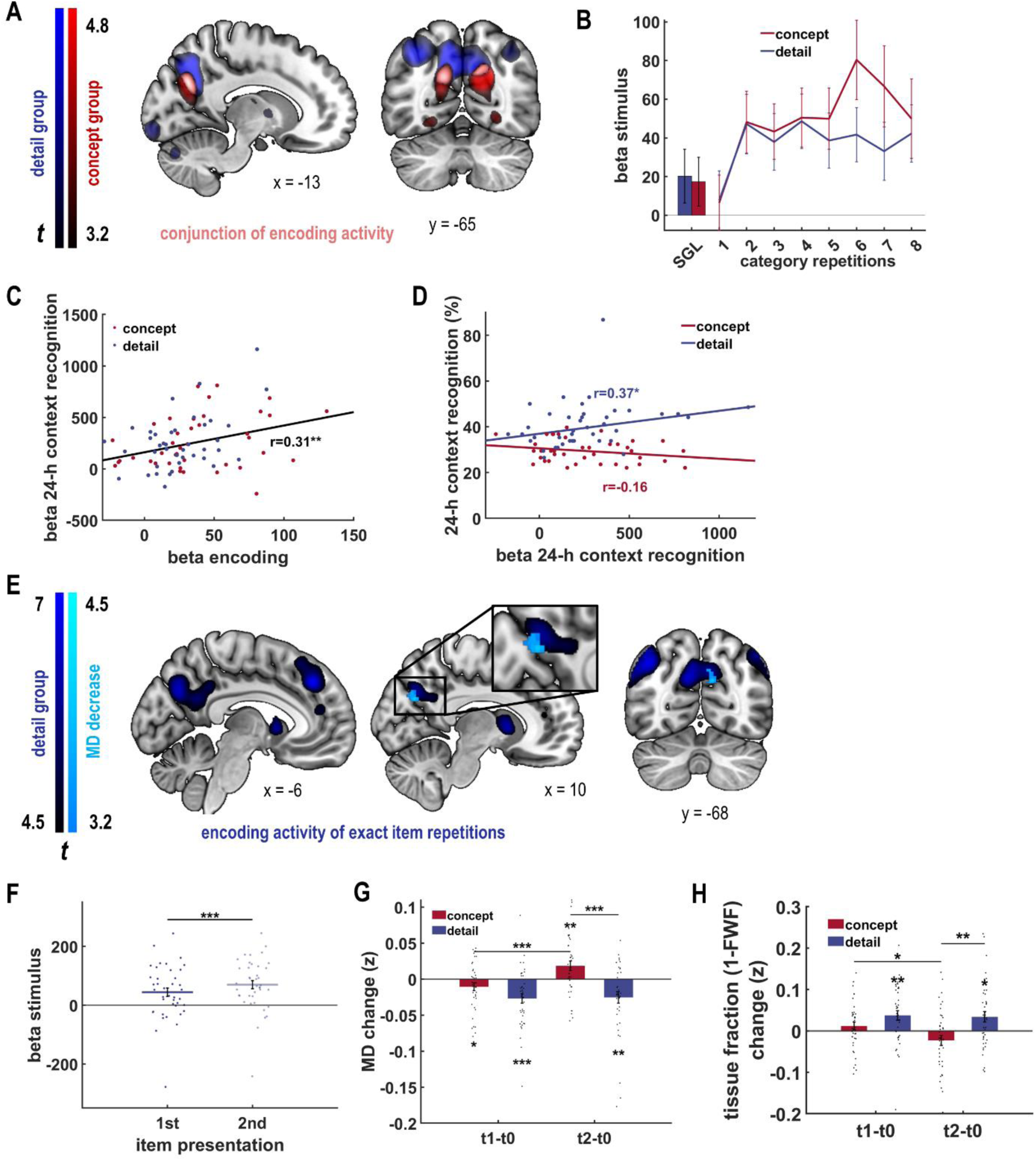
A hub for episodic memory formation in the precuneus. (**A**) During encoding, increased activity in the mPPC in response to conceptual repetitions occurred in both groups (repeated>first category item; pink, conjunction of detail & concept group; for visualization purpose *p*_uncorr_<0.001; peaks are full-volume corrected *p*_FWE_<0.05). While there was an overlap of activity between both groups, group-specific clusters extended to varying degrees into occipital (concept group) and parietal (detail group) cortex (table S5B, C). (**B**) The activity in the conjunction cluster from (A) increased from the first to the second category repetition in both groups and remained stable for all following repetitions (1^st^ compared with 2^nd^ repetition: *t*_78_=-8.24, *p*<0.001; 2^nd^ compared with mean of all following repetitions: *t*_78_=-0.22, *p*=0.83). (**C**) This repetition-sensitive encoding activity predicted activity during stimulus presentation in the recognition task in both groups indicating a reactivation of encoding activity after 24 h. (**D**) Context recognition activity was in turn associated with better context memory in the detail group. (**E** and **F**) The detail group showed increased encoding activity in the mPPC not only to items from the same category but also to exact item repetitions (2^nd^>1^st^ item presentation, *p*_FWE_<0.05, dark blue). This item repetition-sensitive cluster overlapped with the detail learning-specific microstructural changes at t2, after 24 h (t2<t0 × detail>concept, *p*_uncorr_<0.001, k>10, light blue). (**G**) After 24 h, the detail group showed a strong decrease in mean diffusivity (MD) in the precuneus compared with t0 (t2 − t0: *t*_39_=-3.16, *p*=.003) and also compared with the concept group. This decrease was already pronounced after 1 h indicating rapid and persistent microstructural changes that are specific to detail learning. (**H**) The cluster showing detail learning-specific MD changes, also displayed a group-specific increase in the tissue volume fraction (1-FWF) after 24 h and an observable decrease in the detail group already 1h after encoding, indicating a higher relative contribution of brain tissue vs. CSF compartments. Red, concept group; blue, detail group; bars, mean; error bars, SEM; dots, individual data points; *, *p*<0.05; **, *p*<0.01; ***, *p*<0.001.

In a next step, we investigated functional activity changes in response to exact (rather than conceptual) stimulus repetitions in the detail group. In a whole brain analysis, the mPPC displayed a significant increase from the first to the second presentation of the same item (2^nd^>1^st^ item presentation, *p*_FWE_<0.05; Fig. 4E, F, and table S6). Additionally, in a whole-brain dMRI analysis, we observed decreased MD in an overlapping cluster in the mPPC 24 h after encoding in the detail compared with the concept group (t2<t0 × detail>concept, *p*_uncorr_<0.001; Fig. 4E, G and table S4B). This decrease could already be observed at t1 in the detail group, 1 h after encoding (detail group: t1 − t0: *t*_39_=-4.11, *p*<.001, Fig. 4G). Similar to concept learning, the changes in MD are accompanied by a detail-learning specific increase in tissue volume fraction (1-FWF) in the significant cluster after 24h (difference t2 − t0: concept > detail *t*_78_=3.25, *p*=0.002) with the decrease already showing up after 1h in the detail group (t1 − t0: detail group *t*_39_=-3.30, *p*=0.002; concept group *t*_39_=-1.31, *p*=0.20). The analysis of NDI and ODI did not yield any significant group differences (NDI: difference t2 − t0: concept – detail *t*_78_=-1.60, *p*=0.11; ODI: difference t2 − t0: concept – detail *t*_78_=-1.11, *p*=0.27). Taken together, functional and diffusion-weighted results provide further evidence for a rapid emergence of enduring episodic memory-related plasticity in the mPPC.

## Discussion

In the current study, we aimed to better understand rapid neocortical memory formation by investigating the development of conceptual and detailed memory for entirely novel abstract visual stimuli with the help of combined fMRI and dMRI. We found rapid functional and microstructural changes in the neocortex for the two kinds of learning. Learning new visual concepts was accompanied by a linear increase in the functional response and representational changes of early visual cortex. Functional changes persisted for 24 h, were co-localized with microstructural changes, and correlated positively with categorization performance and negatively with context recognition. Learning detailed item-context representations altered the functional response in the medial posterior parietal cortex already at the second encounter of a similar or same item. Furthermore, these functional changes were related to item-context memory performance and overlapped with detail learning-specific changes in microstructure. Thus, we find learning-specific, within-session functional and structural plasticity in the posterior neocortex, suggesting that this region actively participates in memory storage. Our results show that declarative memories can be established rapidly in the neocortex even for information that is unrelated to prior knowledge. Remarkably, also the acquisition of new visual concepts induced rapid plasticity, suggesting within-session formation of new conceptual memories in the neocortex. Unexpectedly, changes occurred very early in the visual processing hierarchy, suggesting that these new abstract concepts are encoded in the same networks that process the corresponding visual features. Furthermore, we extend earlier findings of rapid memory formation of novel associations between existing schemas or concepts in the mPPC (Brodt et al., 2016; Flanagin et al., 2023; Himmer et al., 2019) by showing that this region can rapidly acquire detailed information also in the absence of prior knowledge.

Visual concept formation was localized in early visual cortex showing a gradual increase in functional response across the repeated presentations of each category. Analogous to previously reported episodic-parietal memory formation (Brodt et al., 2016; Gilmore et al., 2015), this increase in early visual cortex activity remained stable over 24 h and predicted successful concept memory. We suggest that the change in occipital activity represents the generalization of shared features across exemplars, and thus the formation of novel visual concepts. In addition to the change in functional activity, we detected persistent group-specific microstructural plasticity in early visual cortex indicating that these concept-learning induced changes are more than a transient processing of conceptual information originating from downstream areas. Instead, they speak for concept memory-related functional and microstructural plasticity in early visual cortex, more specifically in V2 and downstream areas. It might be surprising that an early visual region as V2 shows visual concept memory. However, it is known to handle complex contours, objects, and textures (Hesse and Tsao, 2023; Qiu and von der Heydt, 2005), and even complex category stimuli can be decoded from peripheral regions of V2 and V3 (Vetter et al., 2014). Furthermore, it has been suggested that it stores emotionally salient stimuli (Sacco and Sacchetti, 2010) and plays a critical role in object-recognition memory (Lopez-Aranda et al., 2009). It seems therefore well placed to learn a representation of our stimulus material.

Previous studies of category formation report category selectivity in higher-order visual processing areas and in multimodal regions such as the parietal or prefrontal cortex (Bracci and Op de Beeck, 2016; Fernandino et al., 2022; Jiang et al., 2007; Popham et al., 2021). However, most of these studies used existing concepts (e.g., cars, objects) that needed to be categorized along specific dimensions. Presumably, these existing concepts are already well integrated into semantic networks in higher-order neocortical areas. In contrast, our results suggest that entirely new concepts based on abstract, visual objects can be encoded even earlier in the visual processing hierarchy. Consequently, novel concepts that predominantly depend on perceptual features may be encoded within the same neural circuits that process their corresponding sensory features and provide the necessary level of abstraction (Yee and Thompson-Schill, 2016). In line with this idea, there have been a number of studies indicating concept formation early in the sensory processing streams. Category learning indeed induces category-specific tuning of neuronal responses down to V1 (Ester et al., 2020; Henderson et al., 2025; O’Bryan et al., 2024). Learned category information has even been identified in neurons with retinotopic receptive fields (Goltstein et al., 2021). Similar observations have been made for auditory categories in the auditory cortex (Chillale et al., 2023; Jiang et al., 2018; Xin et al., 2019). Furthermore, two fMRI studies that have investigated category learning of novel objects similar to our current study report functional changes in early visual cortex, although they do not focus on these effects (Gauthier et al., 1999; Hammer et al., 2010).

Our results corroborate those findings, as we not only see a learning-related modification of functional responses in V2, but also distinct changes in representational similarity in V1 with category repetition. The latter finding confirms that even in the earliest visual processing stages, neural responses can be modulated by concept-related information, going beyond simple feature processing (Vetter et al., 2014). The reduction of concept similarity in V1 with increasing category knowledge, upstream of the univariate functional activity increase observed in V2, could be explained by increased top-down modulation in the context of a predictive coding framework (Gilbert and Li, 2013). Here, an emerging concept representation in higher areas would lead to an increased prediction error signal upstream in V1, basically amplifying differences between individual exemplar features and the category representation. Error feedback propagation from higher-order visual processing areas that hold concept information would amplify the unique features of each exemplar that are not part of the concept representation. Similarly, a recent study observed increased perceptual dissimilarity in early visual areas between exemplars and prototypes that reflected the deviation between sensory input and prior expectations (Blank and Bayer, 2022). Together, we show that not only simple perceptual features like grating patterns but also complex categories of abstract visual objects can be represented at an early stage of visual processing in humans. We hypothesize that new, unimodal concepts might initially be processed and encoded in early sensory regions. With more experience they might later travel up along the cortical processing hierarchy while they become increasingly integrated into existing multimodal semantic concepts in parietal and temporal semantic networks.

The gradual activity increase observed during category learning in early visual cortex indicates that the neocortex is able to rapidly abstract new concepts. Remarkably, these changes occurred within a single learning session and thus before a possible reactivation of repeated episodic representations during offline consolidation, which is a prerequisite of semantic memory formation and gist or schema generation in many current theories (de Sousa et al., 2021; Kumaran and McClelland, 2012; Moscovitch et al., 2016; Winocur and Moscovitch, 2011). Thus, our findings support the view that semantic memories can be directly encoded into neocortical networks (Gilboa and Moscovitch, 2021). Importantly, we do not rule out a hippocampal contribution to the rapid emergence of the observed cortical changes but rather suggest that semantic concept formation does not always require slow abstraction from individual episodic memories and could as well develop successively during encoding by rapid extraction of overlapping features directly within the sensory processing streams. The hippocampus might in that case support encoding through a top-down novelty signal (Gomez-Ocadiz et al., 2022). Of note, our task was focused on the memory aspect of category learning, allowing the acquisition of category knowledge across a small number of encounters. Therefore, it is unlikely to reflect the same processes underlying many perceptual category learning tasks often involving either explicit rule-based strategies, implicitly information integration (Ashby and Maddox, 2011; Reinert et al., 2021; Smith et al., 2014) or gist or prototype extraction from individual items or events (Bowman et al., 2020; Gilboa and Moscovitch, 2021; Griffiths et al., 2007; Sekeres et al., 2018). These tasks typically build categories through trial and error across many learning trials and thus emphasize processes of category detection and decision making, like prototype learning (Bowman et al., 2020), information integration, or rule-based strategies (Ashby and Maddox, 2011; Reinert et al., 2021; Smith et al., 2014).

Besides concept learning-specific changes, we also observed changes common to both groups further down the visual processing stream in the mPPC, in particular in the precuneus. This region displayed rapid and stable increases in activity to the repeated presentation of the same or similar items, but it did not increase with concept repetition. In addition, the precuneus displayed a behaviorally relevant increase in activity particularly in response to the exact repetition of an item-context combination in the detail group, and this increase was co-localized with group-specific microstructural plasticity. These results thus add to accumulating evidence that the precuneus represents a hub for episodic memory (Dadario and Sughrue, 2023; Gilmore et al., 2015; Hebscher et al., 2019a; Mazzoni et al., 2019). They extend earlier findings in that not only new associations between existing concepts (Brodt et al., 2018; Brodt et al., 2016; Flanagin et al., 2023; Himmer et al., 2019) but also detailed contextual memories for novel, abstract material are located in mPPC. Surprisingly, the functional changes in response to exact item repetitions were observed even though items were only repeated once, supporting suggestions that the neocortex is capable of one-trial learning (Chindemi et al., 2022; Cichon and Gan, 2015). The particular sensitivity of the posterior neocortex to repeated presentation of information might represent a general mechanism enabling memory for conceptual and episodic information alike, indicating overlapping neural mechanisms for semantic and episodic memory (Gilboa and Moscovitch, 2021; Renoult et al., 2019; Tanguay et al., 2023). It remains for future studies to elucidate how reactivation through learning repetition during wakefulness might represent a different or complementary process to the hippocampus-driven reactivation shown to underlie offline memory consolidation in sleep and wakefulness (Antony et al., 2017; Brodt et al., 2023; Navarro Lobato et al., 2023). In the current study, spontaneous replay during the short offline phases of our encoding task might also have contributed to the rapid consolidation of both detailed and concept memory. However, spontaneous replay or rehearsal is very likely also part of the natural learning process in any learning situation and occurs even within very short inter-stimulus intervals (Halpern et al., 2025). Remarkably, our findings indicate that both the conceptual representations in early visual cortex as well as detailed parietal memory seem to be encoded into the neocortex without the need for offline systems consolidation.

While our study cannot pinpoint the exact mechanism underlying the observed microstructural changes, multi-compartment diffusion modeling with NODDI yielded an increase in the tissue volume fraction for both clusters showing learning-specific changes in MD, a pattern that has previously been reported in the visual cortex after short-term learning of a virtual spatial navigation task (Villemonteix et al., 2023). At the same time, there was no significant change in the relative contribution of the intra- and extracellular tissue compartments, indicating that the observed increase in tissue density might be driven by changes at the level of both neural and glial processes as well as cell somata (Tavor et al., 2013). In line with this reasoning, animal studies have shown that learning-induced decreases in MD after 24 h co-occur with astrocyte activation and corresponding cell swelling but also with an increase in BDNF and a higher number of synaptic vesicles, all potentially indicative of LTP (Blumenfeld-Katzir et al., 2011; Sagi et al., 2012). Similarly, Tavor et al. (2013) observed MD decreases after a spatial navigation task accompanied by an increase in the restricted compartment estimated with the composite hindered and restricted model of diffusion (CHARMED), which can most likely be attributed to remodeling of dendrite and glial processes.

To avoid any link to already existing knowledge structures, task stimuli were designed to rely on combinations of low-level visual features. Categorizing the items in the concept group was based on simple perceptual processes requiring attention to shape and color, which are important features of naturally-occurring basic categories that can be learned without prior knowledge (Rosch et al., 1976). Therefore, our findings might represent a basic mechanism of learning abstractions also in more natural settings. The detail learning task, on the other hand, required attention to fine-grained differences between individual stimuli and arbitrary item-context associations. While the scenes differed from naturalistic situations by the absence of familiarity and the lack of spatial context, episodic memory is always characterized by a uniqueness of the event and a certain arbitrariness of the elements of the scene. Although naturalistic learning will usually have more associations with known concepts and items, we believe that our present task, by focusing on the contrast between novel visual category learning and novel detail learning, elucidates basic mechanisms of learning abstractions and item-context associations. While the presented context was not predictive in the concept group, the consistent co-occurrence of all aliens together with a planet might still have led to inclusion of a general “planetary” context into all learned concepts as a domain-general scene category (Aminoff and Tarr, 2015). However, since our analyses contrasted detail and concept group, and both conditions included planetary context, concept-related functional activity should not be affected by the context.

In conclusion, we provide evidence that the neocortex can rapidly form memory for conceptual as well as for detailed visual information in the absence of prior knowledge or pre-existing schemas. Overall, our results seem more compatible with ideas of direct neocortical or parallel hippocampal/neocortical encoding than with a complementary learning systems model (Cowansage et al., 2014; Gilboa and Moscovitch, 2021). Most importantly, our results indicate that conceptual information does not necessarily rely on a slow process of abstraction from episodic memories. It can emerge directly within the sensory processing stream of the early visual cortex, suggesting that memories are formed within the same neural circuits where the information is processed.

## Materials and Methods

### Participants

81 healthy right-handed participants (44 female, 37 male; age 23.19 ± 3.17 years [M±SD]) were recruited via email. Data from one participant was excluded, as he only showed up on the first experimental day. Experimental procedures were approved by the Ethics Committee of the Eberhard Karls Universität Tübingen, participants gave written informed consent and received financial compensation.

### Procedure

All participants were randomly assigned to either the concept group (n=40) or the detail group (n=40, see section “Encoding task”). The experimental procedure was the same for the two groups (Fig. 1A). Each participant was tested on two consecutive days, exactly 24 hours apart. The first experimental day started with an MR scanning block (diffusion+) including a localizer, a B1 field map and balanced steady state free precession acquisition (bSSFP; not analyzed in this study) and a set of diffusion-weighted images (t0). Afterwards, on the first day of testing, the encoding task was explained and practiced outside of the scanner until the task instructions were understood. Subsequently, the participants performed the encoding task during fMRI (task duration 49.55 ± 6.52 min [M±SD]). Following the task, the participants answered different questionnaires and took a break of 45 min, during which they were expressly forbidden to study. Afterwards, participants underwent a second diffusion+ scan (t1), which was timed so that the second set of diffusion-weighted images was acquired exactly 1 hour (60.00 ± 2.70 min [M±SD]) after the end of the encoding task. Finally, anatomical high-resolution T1 and T2-weighted images were acquired. The second experimental day started with a third diffusion+ scanning block (t2, 23.99 ± 0.13 h after t0) and was followed by a short categorization task outside of the scanner (task duration 1.93 ± 0.36 min [M±SD]). Afterwards, the item-context recognition task was explained and practiced outside of the scanner and performed during fMRI (task duration of 23.4 ± 3.55 min [M±SD]). All tasks performed inside the scanner were projected to a screen at the back of the scanner bore, which the participants could see via a mirror attached to the head coil. All tasks were programmed with Cogent 2000 (v1.33, Wellcome Trust Centre for Neuroimaging, London, UK) and presented via MATLAB R2018b (Mathworks, Sherbom, MA).

### Encoding task

The abstract visual stimuli for the encoding task consisted of an item (“alien”) in the foreground and an image of a planet in the background (context) (Fig. 1B). For creating the items, we used the “growing digital embryos” software (http://vision.psych.umn.edu/users/kersten/kersten-lab/camouflage/digitalembryo.html). More details about the algorithm are provided in Brady and Kersten (2003) (Brady and Kersten, 2003). Starting from geometric volumes, stochastic growth processes and simulated forces are iteratively applied to a simple polyhedron to make it “grow” into more complex polyhedrons that differ in shape. Objects resulting from this process resemble biological growth. Importantly, several similar but different objects can grow out of the same “ancestor”. Ancestors were chosen by the experimenters to be different from each other considering protrusions, overall shape, and orientation in space. Individual stimuli were removed if category features were not present or if similarity to another stimulus from the same category was considered too high. Afterwards, a category-specific colored texture was applied to the stimuli. Textures for each category were created from sections of a blurred picture thus having a similar color palette and similar structure. The background images were based on existing planet images from different openly available websites (representative images in the figures created by Dall-E 3). The items were designed in a way that they either had unique features, so they would be characterized as “single items” or they shared a similar shape and color scheme, so they could be identified as members of the same category (“category items”, Fig. 1B). These categories were designed and piloted to have visually distinct but not fully obvious category-defining features so that category knowledge could be acquired over a few learning trials once the stimuli are known. Unlike typical category learning tasks (e.g., prototype/exemplar learning tasks), our task emphasizes the memory aspect over category detection and decision. The stimulus set was validated in an independent sample with 10 participants (see supplementary methods).

Participants in both learning groups differed only with regard to their encoding goal (Fig. 1B). The detail group was instructed to remember the stimuli in detail, including the unique features of the item as well as the context in which it was presented. The concept group focused on shared features between items to identify item categories.

The background story of the task for the detail group was that the participant was a real estate manager for a group of planets. Each alien (item) was allowed to own exactly one plot of land on each planet. The administrator’s job was to spot those aliens that show up twice on the same planet. To solve the task, the participant had to remember the detailed features of each alien as well as those of the planet (context) it appeared on. The participant was advised that there are aliens that have very similar appearance. For the concept group, the background story was that the participant was a manager in an alien family reunion office. Their task was to find aliens on different planets that belonged to families. To solve the task, the participant had to gradually identify the similarities in shape and color that defined families of aliens (categories). The participant was advised that not all aliens belonged to families but that some were “loners” (single items).

During encoding, both groups were presented with exactly the same stimulus material and made old-new decisions in accordance with their respective encoding goal: The detail group judged whether they had seen the exact item-context combination already before, while the concept group indicated whether they had seen a similar looking item before. Accordingly, they replied either “old” or “new” via a button press of the right or left hand, respectively. They indicated their judgment confidence by either using index or middle finger. Old/new assignments to right and left hand were randomized between the participants.

During encoding, the different visual stimuli were presented successively in 8 fMRI runs with 16 encoding trials each. After each fMRI run, the participant had a break of ∼90 s during which they were instructed to relax and not think about the task. During the break, fMRI was interrupted by a short diffusion-weighted scan (not analyzed in this study). The stimulus set consisted of 32 single items that were only shown once during the whole experiment and 12 categories with 4 items each that were shown twice during the experiment, resulting in a total number of 128 trials. Items belonging to the same category were equally distributed over the 8 fMRI runs. The 16 different planet contexts were pseudo-randomly assigned to the items and equally distributed so that each planet would be shown 8 times in total. 75% of the category items were presented with the same context during their second presentation. To increase distinctiveness and to avoid association of context information to the categories, items of the same category never shared the same context. In a debriefing questionnaire, 39 of the 40 concept learning participants affirmed that they did not attempt to learn any context information. Pairing of items and contexts was randomized between subjects. Each trial started with a fixation cross with a jittered duration of 1.5-2 s followed by the stimulus presentation, which lasted until the participant made a decision via a button press. Feedback was provided by a green (correct) or red (incorrect) frame, and the correct answer (old/new) was displayed below the image for 1.5 s. Additional feedback was provided by 4 s of continued stimulus presentation (for new stimuli), of the same item in front of the original context (for old stimuli, detail group), or of the preceding item of the same category (for old stimuli, concept group). This stimulus was presented in the colored feedback frame. The next trial started after a jittered inter-trial interval (ITI) of 1.5-2 s presenting a fixation cross.

Every ∼80 s, the encoding task was interrupted for ∼15 s by a baseline task, in which participants had to repeatedly judge whether a grey circle was presented sightly right or left to the center of the screen and respond via a button press of their left or right hand. They could further indicate their confidence by either using their index or middle finger. Feedback was provided for 1.5-2 s showing a green plus for correct and a red cross for incorrect responses. Trials were separated by a fixation cross presented for 1 s.

### Categorization task

On the second experimental day participants completed a categorization task, during which participants were presented with novel items that looked similar to items from the encoding task. Both groups received the same task instructions. They were informed that throughout the encoding task, there were aliens belonging to “families” that had resemblance in shape and color as well as “loners” that were unique. We instructed them that we would now present novel items and that they had to judge whether they belonged to a family or resembled one of the unique looking items presented during encoding (doppelgangers). Half of the items were new exemplars from the encoding task categories, the other half resembled one of the unique looking single items. For each item, participants indicated whether the presented item belonged to a previously seen category. The task consisted of 24 trials, in which an item was presented in the center of the screen. Participants indicated yes/no via a button press with the right or left hand, respectively. They could further indicate their confidence by either using their index or middle finger. Yes/no assignments to right and left hand were randomized between the participants. The item was presented until response and was immediately followed by feedback showing a green plus for correct and a red cross for incorrect responses for 1 s.

### Item-Context recognition task

The item-context recognition task was conducted with concurrent fMRI. Each trial consisted of one or two phases. In the first phase, participants were presented with an item and had to judge whether this was an old item that they remember from the encoding task or a new item. For old items, this decision was followed by an alternate forced choice task in which participants had to recognize the context associated with the item out of four options. During encoding, the detail group was instructed that the “correct” context for items presented twice was always the first, initially presented, context. However, in the recognition task, we did not test items that were presented with two different contexts during encoding. The task consisted of 116 trials, with 36 old category items, 32 old single items, 24 new category exemplars and 24 entirely new items. A trial started with the presentation of an item in the center of the screen. Participants indicated old/new with either right or left hand, randomized between the participants. The item was presented until response and was immediately followed by feedback showing a green plus for correct and a red cross for incorrect responses for 800 ms. If the trial showed a new item the feedback was followed by a jittered ITI of 1.5-2 s showing a fixation cross until the next trial started. For old items, the feedback was followed by the presentation of the same item in front of four different contexts each in one corner of the screen. Participants indicated the correct context via button press with their right and left index and middle finger. The three lure contexts were pseudo-randomly selected from the other contexts shown during encoding. The stimuli were presented until response and were immediately followed by feedback and ITI. The recognition task was interrupted every ∼80 s by an odd-even baseline task that lasted for ∼15 s.

### Behavioral data and performance measures

All analyses of behavioral data were conducted in MATALB and were based on the percentage of correct responses during the respective task. To control for response bias, data from the categorization and the item recognition task are reported as d prime (d’), by normalizing the difference of the hit-rate minus false-alarm rate. Statistical testing of performance (categorization, item recognition, and context recognition) relied on two-sample t-tests to compare the memory performance between the two groups. To assess performance changes across the encoding task, a regression analysis of mean accuracy over the 8 encoding runs was performed for each group and the slope was compared between groups in a two-sample t-test. All tests were two-tailed with an α-level of 0.05. To calculate correlations between functional imaging data and memory performance, beta values for the respective contrasts were extracted and Spearman’s rank correlations were calculated. Significance of group differences was determined on Fisher’s z-transformed correlation values.

### MRI data acquisition and analysis

All MRI data was obtained with a 3T Magnetom Prisma scanner (Siemens, Erlangen, Germany) with a standard 64-channel head volume coil at the Max-Planck-Institute for Biological Cybernetics in Tübingen. Functional and diffusion-weighted images were acquired using multi-band sequences provided by the University of Minnesota Center for Magnetic Resonance Research. fMRI BOLD imaging was conducted with an echoplanar imaging (EPI) sequence: 2000 ms repetition time (TR); 30 ms echo time (TE); 72° flip angle (FA); 208×208 mm^2^ field-of-view (FOV); 104×104 nominal matrix size; 2.3 mm slice thickness; 2.0×2.0×2.3 mm^3^ voxel size; 60 transversal slices with interleaved slice acquisition; parallel acquisition technique (PAT) with in-plane acceleration factor 2 (GRAPPA) and multi-band acceleration factor (MB) 2. Additionally, a magnetic B0 field map was acquired for distortion correction of the functional images. For each timepoint, diffusion-weighted images were obtained with a multi-shell acquisition (cmrr_mbep2d_diff, 3500ms TR; 62ms TE; 90° FA; 96 whole-sphere gradient directions distributed between two b-values: 1000 s/mm^2^ (32 directions), 2500 s/mm^2^ (64 directions); 5 interspersed b0 images; 220×220 mm^2^ FOV; 110×110 nominal matrix size; 2 mm slice thickness; 2 mm isotropic voxel size; 70 transversal slices with interleaved slice acquisition; PAT 2 GRAPPA; MB 2), acquired twice with opposite phase encoding directions (AP/PA). For the calculation of MD only the b=1000s/mm^2^ shell was used, while the multi-compartment model NODDI was fit to the full set of images. Additionally, we acquired a B1 map, bSSFP data and a second diffusion protocol at the same time points, and a reduced set of dMRI images throughout the encoding task, which are not analyzed in the current study.

In a separate scanning block, we acquired a high-resolution 3D magnetization-prepared 2 rapid gradient echoes (MP2RAGE) sequence (5000 ms TR; 2.98ms TE; 700/2500 ms inversion time (TI) 1/2; 4°/5° FA 1/2; 256×256×176 mm³ FOV; 256×256×176 nominal matrix size; 1 mm isotropic voxel size; PAT GRAPPA 3×1). In addition, a high-resolution T2-weighted image was acquired with a 3D turbo spin echo (TSE) sampling perfection with application optimized contrasts using different flip angle evolution (SPACE) sequence (3200 ms TR; 564 ms TE; 256×256×166 mm³ FOV; 320×320×208 nominal matrix size; 0.8 mm isotropic voxel size; PAT GRAPPA 2×1).

Where applicable, preprocessing followed the minimal preprocessing pipeline for the Human Connectome Project (Glasser et al., 2013) with the FMRIB Software Library (FSL, v6.0.5) (Jenkinson et al., 2012).

### Anatomical data

After removing background noise from the MP2RAGE-derived T1-weighted “UNI” image as first proposed by O’Brien et al. (https://github.com/JosePMarques/MP2RAGE-related-scripts) (O’Brien et al., 2014), initial brain extraction was done using FreeSurfer’s recon-all (v.7.2.0, http://surfer.nmr.mgh.harvard.edu), subsequently applying the skullstrip gcut option and/or manually editing, if required. The brain-extracted T1-weighted image was preprocessed using fMRIPrep 21.0.2 (Esteban et al., 2019), which is based on Nipype 1.6.1 (Gorgolewski et al., 2011) in the following steps described in boilerplate provided by the software: The T1-weighted (T1w) image was corrected for intensity non-uniformity (INU) with N4BiasFieldCorrection (Tustison et al., 2010), distributed with ANTs 2.3.3 (Avants et al., 2008), and used as T1w-reference throughout the workflow. The T1w-reference was then skull-stripped with a Nipype implementation of the antsBrainExtraction.sh workflow (from ANTs), using OASIS30ANTs as target template. Brain tissue segmentation of cerebrospinal fluid (CSF), white-matter (WM) and gray-matter (GM) was performed on the brain-extracted T1w using FSL fast (Zhang et al., 2001). Brain surfaces were reconstructed using FreeSurfer recon-all (Dale et al., 1999), and the brain mask estimated previously was refined with a custom variation of the method to reconcile ANTs-derived and FreeSurfer-derived segmentations of the cortical gray-matter of Mindboggle (Klein et al., 2017). Volume-based spatial normalization to standard space (MNI152NLin6Asym) was performed through nonlinear registration with antsRegistration (ANTs 2.3.3), using brain-extracted versions of both T1w reference and the T1w template. The following template was selected for spatial normalization: FSL’s MNI ICBM 152 non-linear 6th Generation Asymmetric Average Brain Stereotaxic Registration Model (Evans et al., 2012).

### Functional imaging data

The first five functional volumes of each fMRI run were discarded from analysis to control for magnetic saturation effects. For the encoding task, all remaining volumes of the eight runs were concatenated to one timeseries. After visual inspection of the raw fMRI data, ∼66% of subjects showed artifacts in a small fraction of volumes during either encoding or recognition, or both of the tasks (0.5 ± 1.0% of volumes (encoding), 0.8 ± 1.9% of volumes (recognition) [M±SD]). All affected volumes were replaced with the mean of the preceding and following volume. Subsequently, all volumes were motion corrected using a 6 df mcflirt registration to the mean volume of the whole timeseries. Using epi_reg including the respective B0 field map for distortion correction, the single-band reference of the first run was registered to the individual high-resolution T1-image. Subsequently, all transforms were concatenated to a single warp image for resampling the data to 2 mm MNI space in one single step using antsApplyTransforms (ANTs 2.4.2). Data was intensity normalized to a brain mean of 10000, smoothed with a Gaussian kernel of 6 mm FWHM and high-pass filtered for frequencies higher than 0.01 Hz.

For the univariate analysis, the functional data were analyzed in a two-level model using the SPM12 toolbox (Wellcome Department of Cognitive Neurology, London, UK) in MATLAB. Encoding task: For the fixed effects (first level), individual models for each subject were implemented. Time for encoding and retrieval was not limited in order to allow thorough processing in the beginning and avoid idle periods later in the task. This also enabled the measurement of response time for each trial. We observed the expected decrease in reaction times over the course of the experiment (mean slope=-0.11, t_78_=-6.68, p<.001; no difference between groups: t_78_=0.34, p=0.74). To take this decrease into account and to ensure our findings are independent of presentation time, all first level models were run on the residuals of a first level model that included a regressor with all stimulus onsets parametrically modulated by the reaction time (RT) of each trial. This hierarchical approach assures that no RT-related variance remained in the subsequent model, irrespective of any possible collinearity between RT and other factors. This model further included six motion regressors and 7 constants indicating the 8 separate MR acquisition runs during encoding. For all conditions of interest in the first level models, event duration was modelled corresponding to the respective reaction time. Besides the respective conditions of interest, left and right-hand responses, feedback stimulus, positive and negative task feedback, baseline stimulus, positive and negative baseline feedback were all modelled as separate regressors. The random effects (second level) models were full-factorial with one repeated factor for stimulus type and one between group factor. All models were based on first-level contrast images with the respective baseline activation subtracted from the condition of interest. All contrast images were smoothed with a Gaussian kernel of 8 mm FWHM.

To investigate the formation of a new visual concept, we compared the change of the neural response with increasing category evidence between the two groups modelling a linearly increasing contrast over the eight category repetitions. The fixed effects model included 9 regressors of interest: one regressor with all single items, and one regressor for each of the eight category repetitions. The second level model included one repeated factor with 9 levels and the between group factor. For the interaction between the two groups, we contrasted the linear increase over category repetitions of one group with the corresponding linear decrease of the opposite group. To identify neural activity related to repetition of category information, two first-level regressors were modeled, one consisting of all single items and first items of each category, the other consisting of all following category items. On the second level, a region-of-interest analysis based on the significant cluster from the linear category increase model was performed, testing the betas of repeated category items and single/first category items and their interaction with the group factor to identify group differences in response to familiar and novel items. To investigate the neural response to perceptual repetitions, we looked at functional activity only within the detail group, as exact item repetitions were confounded with category information in the concept group. The corresponding first level model included four separate regressors with onsets of the different stimulus types: single items and first items of each category, the first presentation of all remaining category items, the second presentation of category items with the original context, and the second presentation of category items with a different context. The second level included one repeated factor with 4 levels. We contrasted the neural response of exact item repetitions with first presentations.

While the paramount goal of our design was to keep the visual stimulation during the fMRI events representing learning constant and to have overall similar task difficulty and performance, the feedback that the two groups received throughout the task differed in the number of trials in which the feedback stimulus was different from the target stimulus. However, variance explained by the feedback stimuli is captured in an independent regressor of no interest in our fMRI analysis including their onsets and durations and therefore does not affect the analysis.

Item-context recognition task: Separate fixed effects models were implemented for the item and for the context recognition phase for each subject. For all conditions of interest, event duration was modelled corresponding to the respective reaction time. The first fixed effects model included four regressors of interest: old category items, old single items, new category items and new single items. Furthermore, baseline, context recognition, left and right key presses, positive and negative feedback were modelled in separate regressors. The corresponding second level random effects model for item recognition was full-factorial with one between group factor and one repeated factor with 4 levels based on first-level contrast images with the respective baseline activation subtracted from the condition of interest.

The second fixed effects model was implemented to model activity related to context recognition and included the following regressors of interest: correct context recognition of single items, incorrect context recognition of single items, correct context recognition of category items, incorrect context recognition of category items. Furthermore, baseline, item recognition, left and right key presses, and positive and negative feedback were modelled in separate regressors. The corresponding second level random effects model for context recognition was full-factorial with one between group factor and two repeated factors (correctness and stimulus type), each with two levels based on first-level contrast images with the respective baseline activation subtracted from the condition of interest.

To identify areas of overlapping activity across the two groups, conjunction analyses were conducted using the conjunction of minimum statistic analyses (Nichols et al., 2005), that takes the maximum p-value of all individual analyses per voxel.

### Diffusion-weighted imaging data

While mean diffusivity (MD) estimated with diffusion tensor imaging reflects non-directional changes in diffusivity independent of the underlying tissue, NODDI constitutes a multi-compartment model assuming three compartments incorporating either unrestricted, hindered or restricted diffusion to reflect the properties of the underlying tissue, namely CSF, extracellular (cell somata) and intracellular space (axons, dendrites). Contribution of the signal from these compartments is estimated in two steps with first identifying the amount of signal coming from unrestricted diffusion within a voxel, reflected by the free water fraction (FWF) and the remaining voxel fraction characterized as the tissue volume fraction (1-FWF). Subsequently, the tissue signal is divided into signal coming from the extracellular (hindered diffusion) and intracellular compartment (restricted diffusion). The neurite density index (NDI) describes the relative volume fraction of the intracellular vs. extracellular compartment. Additionally, the intracellular compartment is characterized by the dispersion of neurite orientations quantified by the orientation dispersion index (ODI) (Daducci et al., 2015).

Within each time point (t0/t1/t2), all diffusion-weighted images were denoised using the Marcenko-Pastur PCA algorithm via DIPY’s mccpa, and motion- and distortion-corrected via a combination of FSL’s topup and eddy. Diffusion tensors were estimated on the b=0 and b=1000s/mm^2^ images with dtifit and based on eddy-rotated b-vectors. NODDI was fit to the full set of denoised, motion- and eddy-corrected diffusion-weighted images using the Accelerated Microstructure Imaging via Convex Optimization (AMICO) framework (Daducci et al., 2015) with the intrinsic parallel diffusivity optimized for parameter estimation in gray matter (d_in_= 1.1×10^−3^ mm^2^/s) (Fukutomi et al., 2018). To improve registration to the anatomical image a shared brain mask for all time points was calculated based on the eddy-corrected b0 images. A linear 7 df registration of the fractional anisotropy (FA) image of each timepoint onto the individual high-resolution T1 scan was calculated with FSL’s flirt and subsequently combined with the non-linear warp to MNI space from fmriprep to resample from undistorted diffusion into MNI space in one single step using antsApplyTransforms (ANTs 2.4.2). MNI space data was smoothed with a Gaussian kernel of 6 mm FWHM to improve inter-subject comparability. For each diffusion parameter (MD, FWF, NDI, ODI), data was z-standardized with respect to the whole gray matter mean separately for each subject and time point to exclude session effects.

To identify brain regions showing differences in microstructural changes between the two groups whole-brain analyses on MD data in MNI space were conducted. Statistical testing was performed with FSL and relied on two 2 (time) × 2 (group) factorial models, comparing MD values of the first (t1, 1 h after encoding), or the second timepoint (t2, 24 h after baseline), respectively, to baseline (t0) and between the two groups. To assess the pattern underlying the significant interaction and to verify that there was an actual decrease of MD in one of the two groups, the difference of the cluster-median of the z-standardized MD-values extracted from the significant clusters was subsequently compared between the two groups with two-sided post-hoc t-tests. Statistical inferences on whole-brain diffusion data are uncorrected for multiple comparisons but only clusters exceeding 10 voxels are reported. Based on the clusters showing significant changes in MD, ROI analyses for the three indices estimated with NODDI were conducted to characterize the observed changes in MD with regard to the underlying microstructure. For FWF-values the difference of the cluster-median of the z-standardized values were compared between the two groups with two-sided post-hoc t-tests, while for NDI and ODI, tissue-weighted means of the z-standardized values from the ROI were calculated first, correcting for partial volume effects of free water (Parker et al., 2021). The tissue-weighted means were then compared between the two groups with two-sided post-hoc t-tests.

### Pattern similarity analysis

To investigate the change in pattern similarity of individual category members to the mean category pattern, the parametrically modulated (see paragraph “functional imaging data”) time-series of encoding activity was used to set up fixed effects first level models to estimate single-trial activity of all category items. We used the least squares single approach (Mumford et al., 2014) to estimate each trial’s activation in a separate model with one regressor of interest including the onset of the respective trial, and two other regressors including all remaining category and single item trials respectively. All models included the same regressors of no interest as the models for estimating univariate activity (see paragraph “functional imaging data”). Trial-specific beta images were used to calculate pattern similarity between each of the eight category item repetitions and the mean of all remaining items of the same category in a whole-brain spherical searchlight analysis with a radius of three voxels. To remove activity unrelated to the respective category, for each individual item the average pattern of all other items from the same repetition was subtracted before correlation. The individual item-category correlation patterns were Fisher’s *z*-transformed and averaged across the twelve categories for each of the eight sequential category presentations. To ensure that the changes in similarity were specific to each category, we additionally calculated the change in pattern similarity over category repetitions of each category item to the mean category patterns of all other categories. There was no change in the similarity of individual items to mean category pattern of different categories (*p*_FWE_>0.15 for all brain voxels). To characterize the changes in pattern similarity of individual items to the mean pattern of the respective category, we compared the linear change of pattern similarity (slope) over category repetitions in a linear mixed effects model including linear increase and group membership as factors to compare the change of pattern similarity between the two groups. Comparisons were conducted for each voxel and full-volume FWE-corrected with *p*<0.05.

## Supporting information

Dataset Encoding Task

## Acknowledgments

None.

## Author contributions

SK, SG, SB wrote the manuscript

KS, VK contributed to editing the manuscript

BF, ME, KS, SG, SB designed the research

SK, AS, VK, PI, SS, FV, SG, SB conducted the experiments; SK, BF, PI, SB analyzed the behavioral data

SK, SG, SB analyzed the neuroimaging data

