## Supplementary Material for "Direct encoding of new visual concepts in early visual cortex"

Svenja Klinkowski *et al.*

**This PDF file includes:**

Supplementary Text
Figs. S1 to S3
Tables S1 to S6

**Other Supplementary Materials for this manuscript include the following:**
Data S1

### Supplementary Text

Encoding Task

In order to disentangle learning and time, categories did not all start in the first run but were distributed over the course of the task. The distribution of category repetitions over the course of the encoding task is visualized in Fig. S1A, showing one exemplary subject. The dotted vertical lines indicate the end of each fMRI run. The plot shows that even in the 5th run a new category was introduced and the second to last repetition of the first two categories occurred already in the 4th run before the last category was introduced. Similarly, first and second repetitions of the same item were distributed across the duration of the entire encoding task (Fig. S1B). Even in the first run, some items were already repeated, and new items were still introduced during the last run of the experiment.

Validation of stimulus set

The perceived similarity of all items from encoding and categorization was validated in an independent sample (n=10, 5 female). For this experiment, we used the so-called “Multiple Arrangement Task” provided by Meadows (https://meadows-research.com/) following the idea of inverse multidimensional scaling[70]. We set a fixed time of 60 min for the participant’s assessment. Participants were instructed to sort the items by similarity based on shape and color. They could drag and drop items in a designated screen area (“arena”) and arrange them according to their similarity with distance reflecting dissimilarity. In the first trial, the participants were presented with all items at the same time for a first broad assessment. In the following trials, different subsamples of the items were presented for a more fine-grained assessment. During the 60 min, the participants completed an average of 50±36 [M±SD] trials. Based on the arrangements of all trials an integrated representational dissimilarity matrix (RDM) was calculated with Euclidean distances indicating pairwise similarities for all assessed items.

To validate higher within-category similarity than between-category similarity, as well as similar distances between all single items and different categories, we compared the mean distances of the corresponding combinations of different items. For all comparisons, we first calculated a within-participant mean of the corresponding item combinations and then compared the condition-specific means within all participants in a paired sample t-test. Members of the same category were rated to be significantly more similar to each other than to members of other categories (mean distances within-category=0.0038, between-category=0.0125; $t_9$=-14.69, $p$<0.001) validating that our predefined categories where perceived as such. Category items of different categories were rated to be as similar to each other as different single items to each other (mean distances between-category=0.0125, between-singles=0.0128; $t_9$=-0.51, $p$=0.52) showing that the different categories did not have a higher similarity to each other than the single items. Furthermore, to assess, whether the new category members presented in the categorization task represent “typical” items for this category, we compare the average distance of all “old” category members to each other to the mean distance of the new exemplar to each of the old ones. The new category members did not differ from the other category members in their similarity (mean distances within-category old exemplars: 0.0038, new exemplar to old exemplars: 0.0037; $t_9$=0.29, $p$=0.78). Similarly, we confirmed that the “doppelgangers” of the single items in the categorization task were more similar to their corresponding items than all other singles to each other (mean distance within-doppelgangers: 0.0024, between-singles:

0.0128; $t_9$=-14.38, $p$<0.001). The average representational dissimilarity matrix (RDM) of all participants is displayed in figure S2A including the four category members shown during encoding and the fifth category member shown during categorization of each of the 12 categories, followed by 12 single items that were shown during encoding and their corresponding doppelganger from the categorization task, and the remaining 24 single items that were only presented during encoding.

In another approach to compare within-category similarity with between-category similarity, we correlated shapes and color histograms of all category items, doppelganger items, and single items from encoding and the categorization task. Average color histogram correlations: r=0.73 (within-category), r=0.21 (between-category), r=0.75 (within doppelganger items), r=0.24 (between single items). Average shape correlations: r=0.51 (within-category), r=0.12 (between-category), r=0.59 (within doppelganger items), r=0.25 (between single items). Although relatively crude measures of obvious similarity only, results show that category items and doppelgangers are visibly more similar to each other than to other items. Summed, weighted correlations with shape and color can be found in figure S2B.

Supplementary Results

During item recognition, the response pattern of the two groups differed depending on stimulus type (Fig. S3A, B). Category items were more likely to be correctly recognized as “old” (hit), while single items were more likely to be recognized as “new” (CR). This response pattern can be explained by the nature of the encoding task, as single items have only been presented once during encoding and are less likely to be remembered (Fig. S3C). Interestingly, the bias to recognize a category item as “old” is stronger in the concept group that explicitly encoded categories, which might be a result of stronger generalization of the category concept. To account for this bias, we reported the bias-adjusted overall performance as d’ in the main manuscript, which does not differ between the two groups. For correctly remembered items (hits), context recognition was 37.7% (±11.4%) [concept group: 31.3% (±7.2%); detail group: 44.0% (±11.4%)], while context recognition for forgotten items (misses) was 30.1% (±11.3%) [concept group: 25.6% (±8.6%); detail group: 34.6% (±12.0%)]. Thus, context memory was higher after correct item recognition in both groups ($F_{1,78}$=26.13, $p$<0.001), however the concept group shows context memory above the chance level of 25% only for recognized ($t_{39}$=5.54, $p$<0.001) and not for forgotten items ($t_{39}$=0.46, $p$=0.65, Fig. S3D).

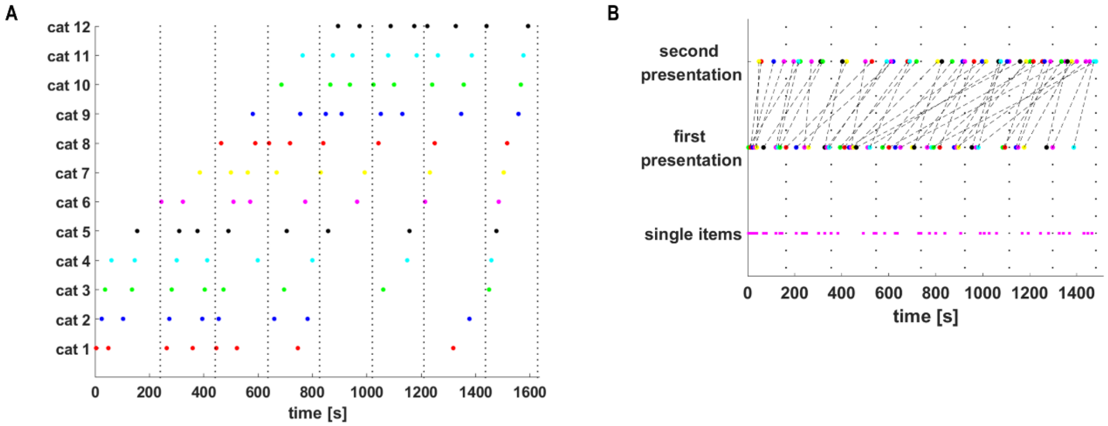

**Fig. S1. Exemplary distribution of onsets.**

(**A**) Category repetitions for each category ("cat 1-12") were distributed over the course of the whole task independent of the 8 fMRI runs (dotted vertical lines). Dots of the same color indicate repetitions of the same category. (**B**) Exact repetitions of all items were distributed across the course of the task with some items being introduced and repeated earlier and others later in the task. Single items are not repeated but also distributed across the whole task. Dots of the same color indicate same items for the repeated category items. Dashed black lines link the first and second presentation of the same item. Single items are plotted in pink. (A and B) Vertical dotted black lines indicate the end of an fMRI run.

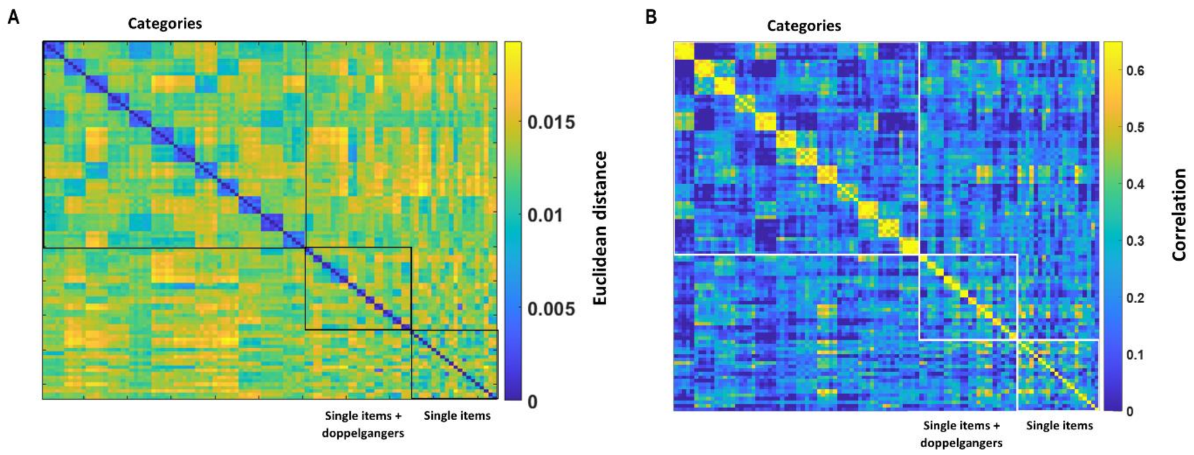

**Fig. S2. Assessment of similarity between experimental stimuli.**

(**A**) To validate the stimulus set, perceived similarity of all category items and single items from encoding and their corresponding doppelgangers and new category items from the categorization task were assessed in an independent sample. The averaged Euclidean distances between all assessed items are displayed in a representational dissimilarity matrix. (**B**) Perceptual similarity of the same items was further validated in a mathematical approach separately correlating shape and color features between all items. The figure shows summed, weighted correlations between all assessed stimuli.

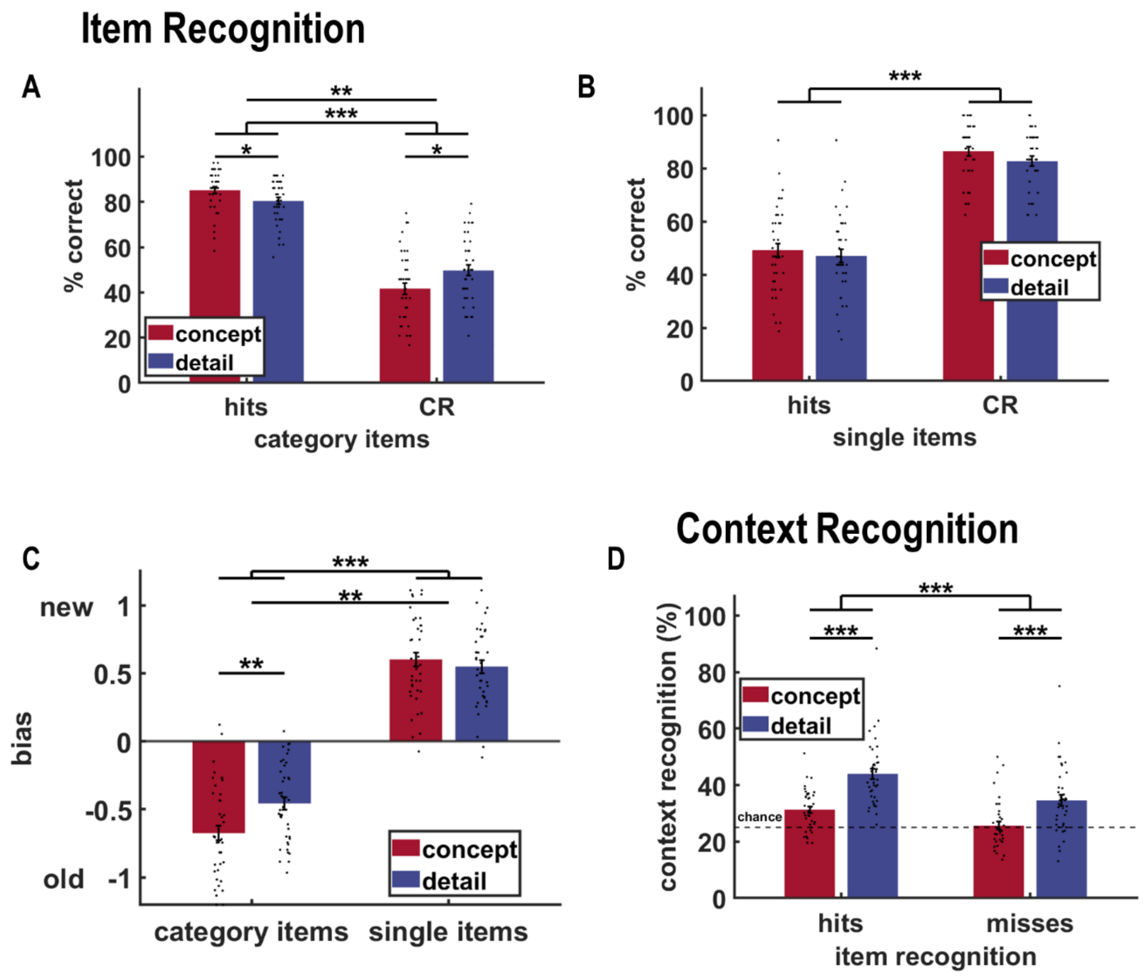

**Fig. S3. Supplementary behavioral results.**

**(A** and **B)** During item recognition on the second day of testing, the two groups differed in their pattern of response to category items ($F_{1,78}$=7.77, $p$=0.007). While both groups were more likely to correctly recognize old category items as "old" (hits) than to correctly rejecting new category items (main effect trial type: $F_{1,78}$=263.42, $p$<0.001), this pattern was stronger in the concept group (hits: $t_{78}$=2.24, $p$=0.03; CR: $t_{78}$=-2.31, $p$=0.02). For single items, the opposite pattern occurs: both groups are more likely to miss old single items and correctly reject new single items (main effect trial type: $F_{1,78}$=275.27, $p$<0.001). **(C)** The different response patterns during item recognition result in a different response bias to category and single items (main effect stimulus type: $F_{1,78}$=705.62, $p$<0.001) with a bias towards responding "old" for category items and a bias towards "new" for single items in both groups. The two groups differ in the magnitude of their bias ($F_{1,78}$=9.94, $p$=0.002) with a stronger bias towards "old" for category items in the concept group ($t_{78}$=-3.09, $p$=0.003) speaking for a higher generalization of the category concept to new (similar looking) category items in the group that focused on similarities during encoding. **(D)** After correctly recognizing "old" items during item recognition (hits), both groups were more likely to remember the corresponding context (main effect trial type: $F_{1,78}$=26.13, $p$<0.001).

**Table S1. Change in encoding task functional activity over category repetitions.**

| Region | $k_{voxels}$ | $p_{cluster}$ | $X_{peak}$ | $Y_{peak}$ | $Z_{peak}$ | $p_{FWE}$ | $T_{peak}$ |
|---|---|---|---|---|---|---|---|
| **Increase over category repetitions × concept>detail** | | | | | | | |
| Right Lingual Gyrus (V2) | 4774 | <0.001 | 22 | -56 | 6 | <0.001 | 7.38 |
| Right Lingual Gyrus (V2) | | | 10 | -68 | -2 | <0.001 | 6.62 |
| Left Lingual Gyrus (V2) | | | -4 | -88 | 22 | <0.001 | 6.62 |
| Supplementary Motor Area (pre-SMA) | 543 | <0.001 | -4 | 2 | 56 | <0.001 | 6.57 |

Regions showing a significantly higher increase in activity over category repetitions in the concept group compared to the detail group. The opposite contrast does not yield significance in any region. The regions listed above exhibited significant peak voxel effects at full-volume family-wise error (FWE) corrected $p_{FWE}<.05$, df=702, and had a cluster size>10. Peak voxel MNI coordinates X, Y, and Z are given in millimeters.

**Table S2. Change in pattern similarity of category items with mean category pattern.**

| Region | $N_{voxels}$ | $X_{peak}$ | $Y_{peak}$ | $Z_{peak}$ | $p_{FWE}$ | $T_{peak}$ |
|---|---|---|---|---|---|---|
| **Decrease of pattern similarity × concept>detail** | | | | | | |
| Occipital Pole (V1) | 3 | -18 | -90 | -4 | 0.026 | 4.83 |

Regions showing a significantly higher change in pattern similarity of individual category items to the mean category pattern over category repetitions in the concept group compared to the detail group. The opposite contrast does not yield significance in any region. The regions listed exhibited significant peak voxel effects at full-volume corrected $p_{FWE}<.05$, df=636. Peak voxel MNI coordinates X, Y, and Z are given in millimeters.

**Table S3. Task-related functional activity 24 h after encoding during item-context recognition.**

| Region | $k_{voxels}$ | $p_{cluster}$ | $X_{peak}$ | $Y_{peak}$ | $Z_{peak}$ | $p_{FWE}$ | $T_{peak}$ |
|---|---|---|---|---|---|---|---|
| **A – Context recognition>baseline × concept>detail** | | | | | | | |
| Right Occipital Fusiform Gyrus (V1) | 13 | 0.020 | 26 | -68 | 0 | 0.020 | 4.78 |
| **B – Context recognition>baseline × detail>concept** | | | | | | | |
| Left Inferior Lateral Occipital Cortex | 48 | 0.006 | -56 | -70 | -4 | 0.021 | 4.77 |
| Left Middle Temporal Gyrus | | | -56 | -56 | -6 | 0.021 | 4.77 |
| Left Middle Frontal Gyrus | 24 | 0.013 | -50 | 34 | 26 | 0.017 | 4.82 |

Regions showing a significantly higher activity during the context recognition task compared to baseline in interaction with learning group. The regions listed exhibited significant peak voxel effects at full-volume corrected $p_{FWE}$<.05, df=312. Peak voxel MNI coordinates X, Y, and Z are given in millimeters.

**Table S4. Long-term learning-induced microstructural changes.**

| Region | $N_{voxels}$ | $X_{peak}$ | $Y_{peak}$ | $Z_{peak}$ | $T_{peak}$ |
|---|---|---|---|---|---|
| **A – Interaction MD decrease (t2<t0) × (concept>detail)** | | | | | |
| Left Intracalcarine Cortex (V1) | 13 | -10 | -78 | 6 | 4.19 |
| Right Precentral Gyrus | 11 | 44 | -2 | 54 | 4.00 |
| **B – Interaction MD decrease (t2<t0) × (detail>concept)** | | | | | |
| Right Precuneus | 50 | 10 | -68 | 28 | 4.18 |
| Left Inferior Temporal Gyrus, temporooccipital part | 48 | -60 | -58 | -16 | 4.24 |
| Left Superior Parietal Lobule | 27 | -32 | -42 | 50 | 4.15 |
| Left Inferior Temporal Gyrus, posterior division | 20 | -48 | -22 | -32 | 3.43 |

Regions showing a significantly stronger decrease in mean diffusivity (MD) from t0 (baseline, before encoding) to t2 (24 h later) in interaction with learning group. The regions listed exhibited significant peak voxel effects at $p_{uncorr} \leq .001$, df=78 and had a cluster size>10. Peak voxel MNI coordinates X, Y and Z are given in mm.

**Table S5. Increased encoding activity in response to conceptual item repetitions.**

| Region | $k_{voxels}$ | $p_{cluster}$ | $X_{peak}$ | $Y_{peak}$ | $Z_{peak}$ | $p_{FWE}$ | $T_{peak}$ |
|---|---|---|---|---|---|---|---|
| **A – Repeated>first, conjunction of concept & detail group** | | | | | | | |
| Right Precuneus | 52 | 0.008 | 18 | -60 | 32 | 0.004 | 5.20 |
| Left Precuneus | 26 | 0.016 | -14 | -68 | 30 | 0.008 | 5.05 |
| **B – Repeated>first, main effect in concept group** | | | | | | | |
| Right Precuneus | 424 | <0.001 | 18 | -62 | 28 | <0.001 | 7.02 |
| Right Precuneus | | | 20 | -56 | 14 | 0.002 | 5.43 |
| Left Precuneus | 74 | 0.005 | -16 | -70 | 30 | 0.006 | 5.11 |
| **C – Repeated>first, main effect in detail group** | | | | | | | |
| Left Precuneus | 748 | <0.001 | -8 | -68 | 40 | <0.001 | 6.77 |
| Right Precuneus | | | 18 | -60 | 32 | 0.004 | 5.20 |
| Right Precuneus | | | 8 | -68 | 38 | 0.006 | 5.13 |
| Left Superior Lateral Occipital Cortex | 440 | <0.001 | -36 | -58 | 42 | <0.001 | 6.61 |
| Right Caudate | 165 | 0.001 | 8 | 6 | 8 | <0.001 | 6.33 |
| Superior Frontal Gyrus | 103 | 0.003 | 0 | 42 | 44 | 0.001 | 5.53 |
| Right Superior Parietal Lobule | 127 | 0.002 | 36 | -50 | 40 | 0.004 | 5.21 |
| Left Caudate | 77 | 0.005 | -8 | 2 | 10 | 0.008 | 5.05 |
| Left Insular Cortex | 67 | 0.006 | -28 | 22 | -6 | 0.001 | 5.56 |
| Right Insular Cortex | 30 | 0.014 | 30 | 24 | -4 | 0.009 | 5.02 |

Regions showing a significantly higher activity in response to repeated compared to first category items and single items in (A) a conjunction analysis of both groups and separately for (B) the concept and (C) the detail group. The regions listed exhibited significant peak voxel effects at full-volume corrected $p_{FWE}<.05$, df=156, and had a cluster size>10. Peak voxel MNI coordinates X, Y, and Z are given in millimeters.

**Table S6. Increased encoding activity in response to exact item repetitions.**

| Region | $k_{voxels}$ | $p_{cluster}$ | $X_{peak}$ | $Y_{peak}$ | $Z_{peak}$ | $p_{FWE}$ | $T_{peak}$ |
|---|---|---|---|---|---|---|---|
| **2nd>1st item presentation, main effect in detail group** | | | | | | | |
| Left Superior Lateral Occipital Cortex | 1785 | <0.001 | -42 | -70 | 42 | <0.001 | 6.89 |
| Left Angular Gyrus | | | -54 | -56 | 32 | <0.001 | 5.79 |
| Left Precuneus | 1688 | <0.001 | -8 | -70 | 32 | <0.001 | 6.50 |
| Left Precuneus | | | -12 | -56 | 24 | <0.001 | 6.12 |
| Right Precuneus | | | 8 | -66 | 34 | <0.001 | 5.88 |
| Right Angular Gyrus | 1474 | <0.001 | 58 | -60 | 36 | <0.001 | 6.18 |
| Left Superior Lateral Occipital Cortex | | | 48 | -66 | 48 | <0.001 | 6.10 |
| Left Superior Frontal Gyrus | 1471 | <0.001 | -2 | 40 | 42 | <0.001 | 7.58 |
| Right Frontal Pole | | | 8 | 40 | 56 | <0.001 | 5.67 |
| Right Superior Frontal Gyrus | | | 10 | 26 | 58 | 0.002 | 5.20 |
| Left Middle Frontal Gyrus | 802 | <0.001 | -42 | 20 | 34 | <0.001 | 6.33 |
| Left Middle Frontal Gyrus | | | -50 | 16 | 48 | <0.001 | 6.23 |
| Right Posterior Middle Temporal Gyrus | 423 | <0.001 | 64 | -30 | -8 | <0.001 | 6.63 |
| Left Posterior Middle Temporal Gyrus | 367 | <0.001 | -64 | -34 | -10 | <0.001 | 6.14 |
| Right Middle Frontal Gyrus | 296 | <0.001 | 46 | 16 | 48 | 0.001 | 5.48 |
| Right Middle Frontal Gyrus | | | 50 | 26 | 28 | 0.024 | 4.65 |
| Right Caudate | 276 | <0.001 | 8 | 10 | 4 | <0.001 | 6.42 |
| Left Caudate | | | -6 | 10 | 2 | <0.001 | 6.01 |
| Left Frontal Pole | 215 | 0.001 | -46 | 54 | 0 | 0.001 | 5.32 |
| Right Insular Cortex | 117 | 0.003 | 32 | 24 | -2 | <0.001 | 5.81 |
| Left Frontal Pole | 104 | 0.003 | -18 | 62 | 6 | 0.007 | 4.95 |
| Left Paracingulate Gyrus | 93 | 0.004 | -10 | 42 | 16 | 0.003 | 5.16 |
| Left Insular Cortex | 88 | 0.005 | -30 | 24 | -2 | <0.001 | 5.68 |

Regions showing a significantly higher activity in response to exact item repetitions compared to their first presentations in the detail group. The opposite contrast does not yield significance in any region. The regions listed exhibited significant peak voxel effects at full-volume corrected $p_{FWE}<.05$, df=312. Peak voxel MNI coordinates X, Y, and Z are given in millimeters.

**Data S1. Stimulus set encoding task.**

The data set includes all abstract visual items that were presented during encoding. They consist of 48 category stimuli including 12 categories with each four different category members. Furthermore, it includes 32 unique looking single items.
