## Supplementary figures and images for "Direct encoding of new visual concepts in early visual cortex"

### Dataset Encoding Task

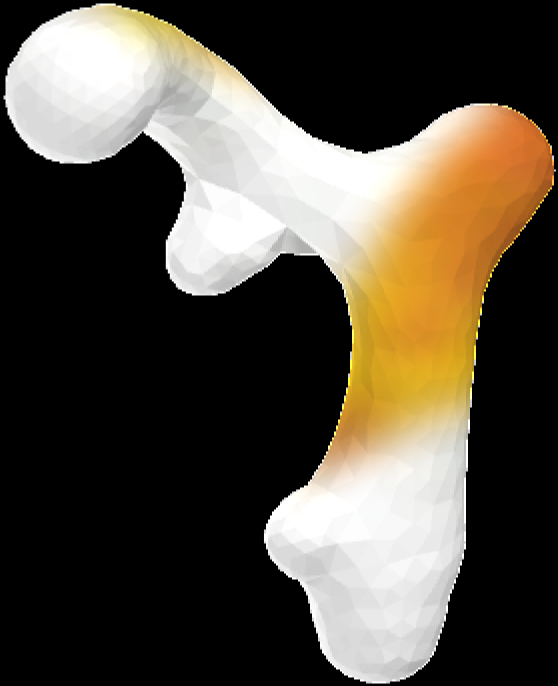

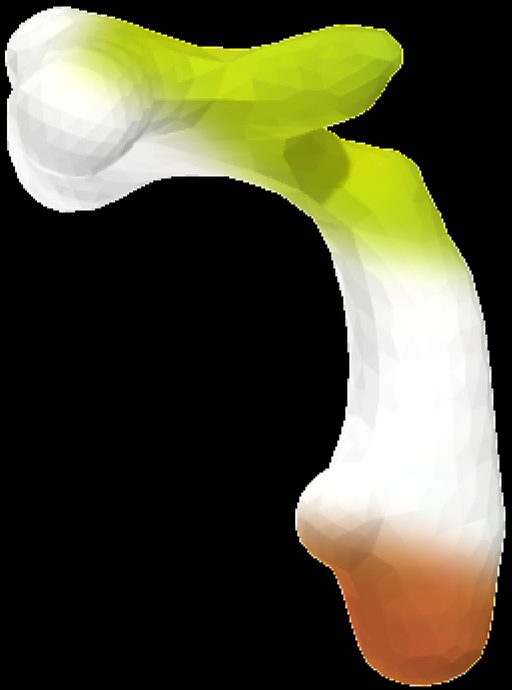

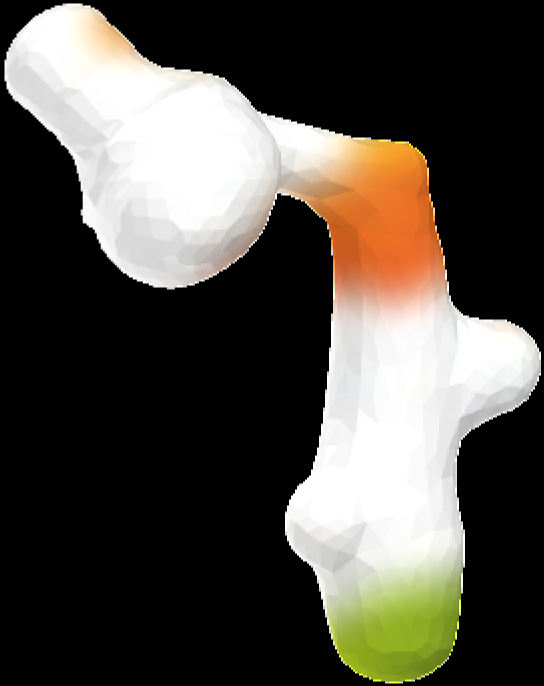

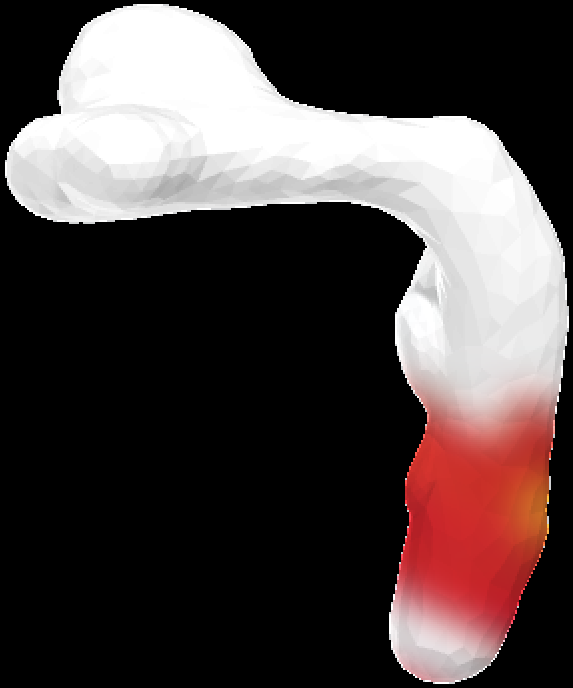

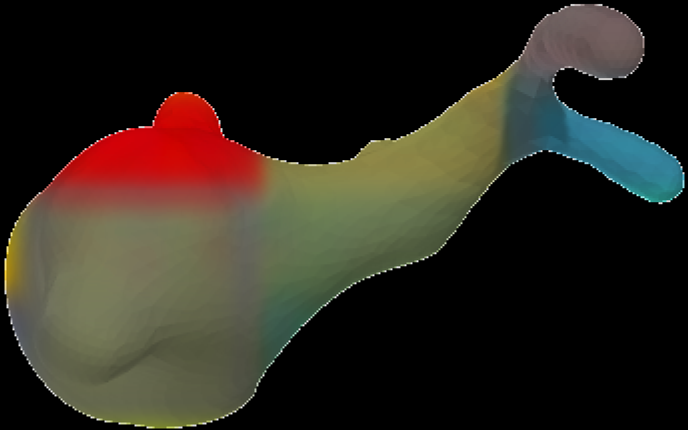

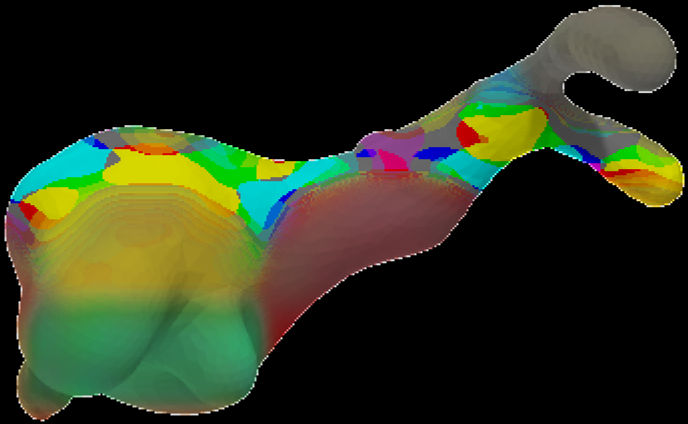

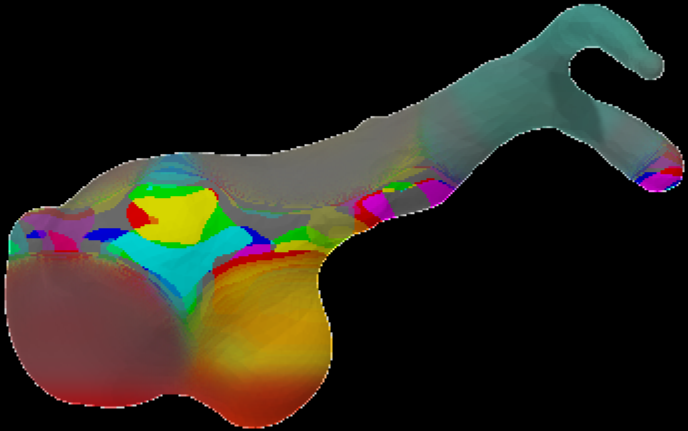

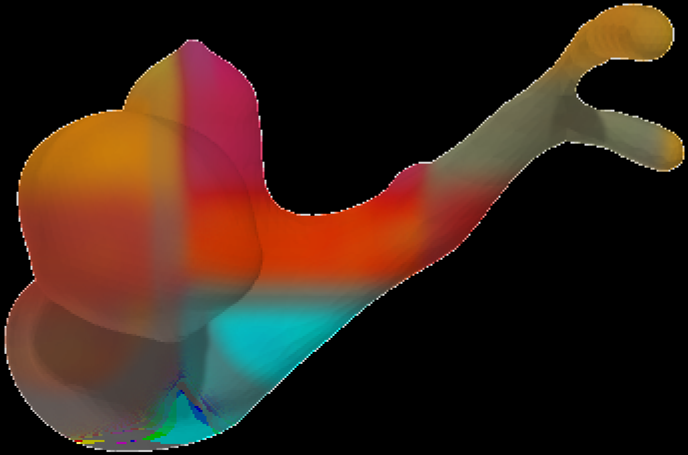

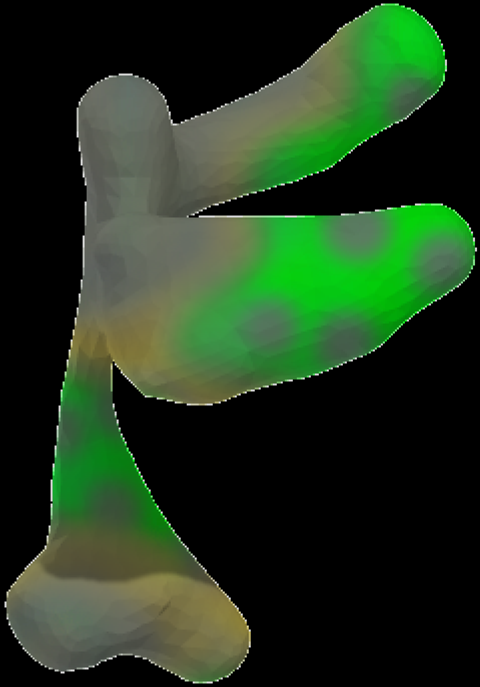

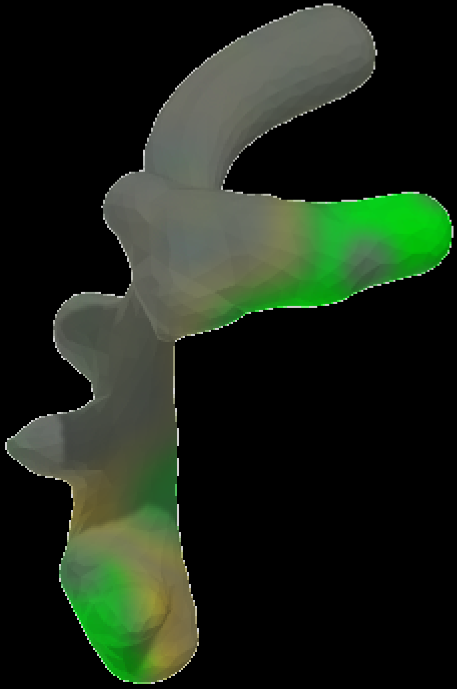

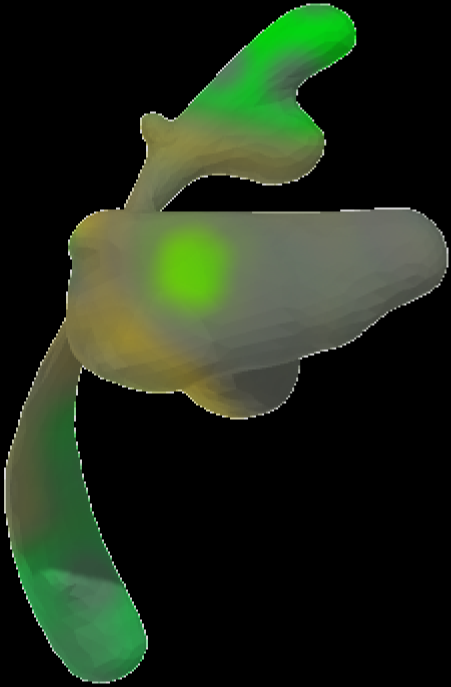

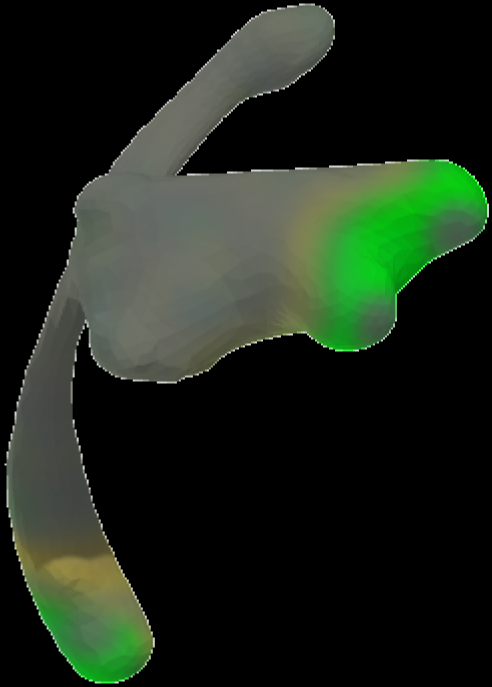

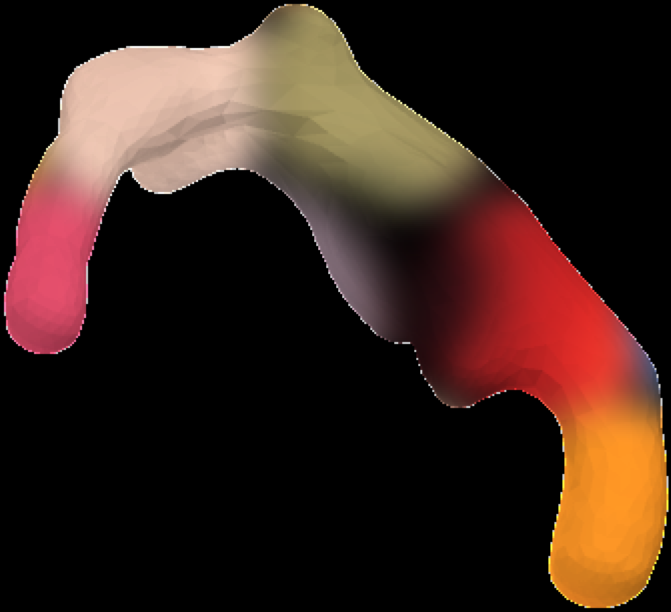

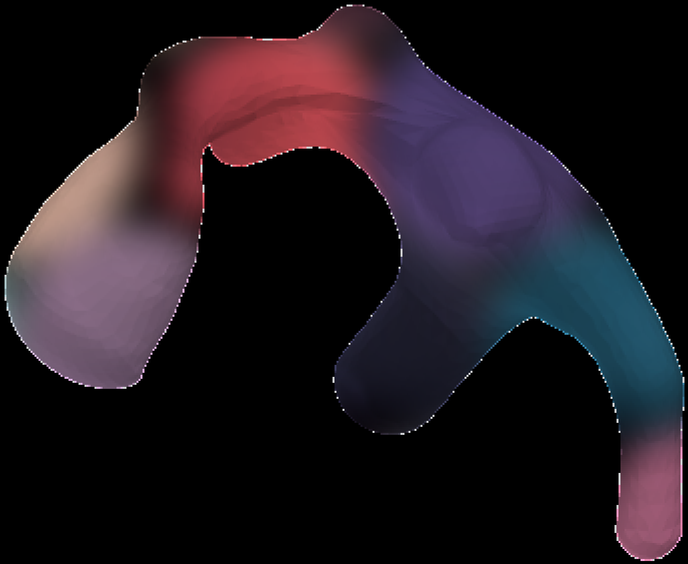

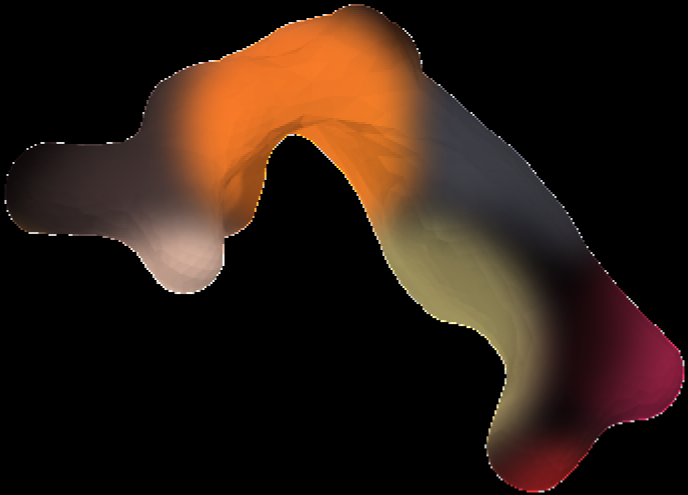

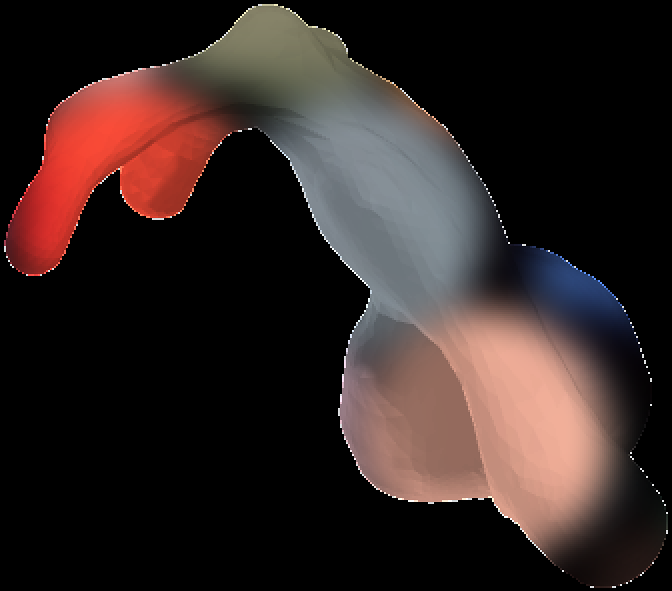

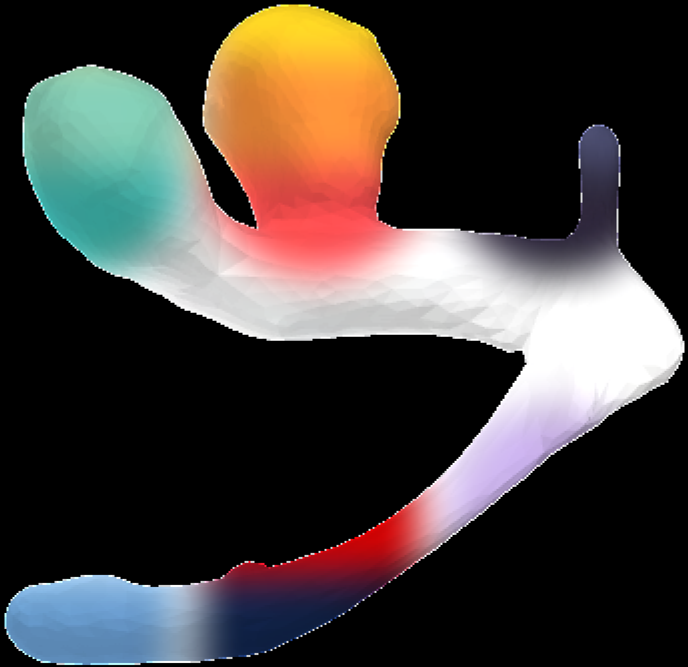

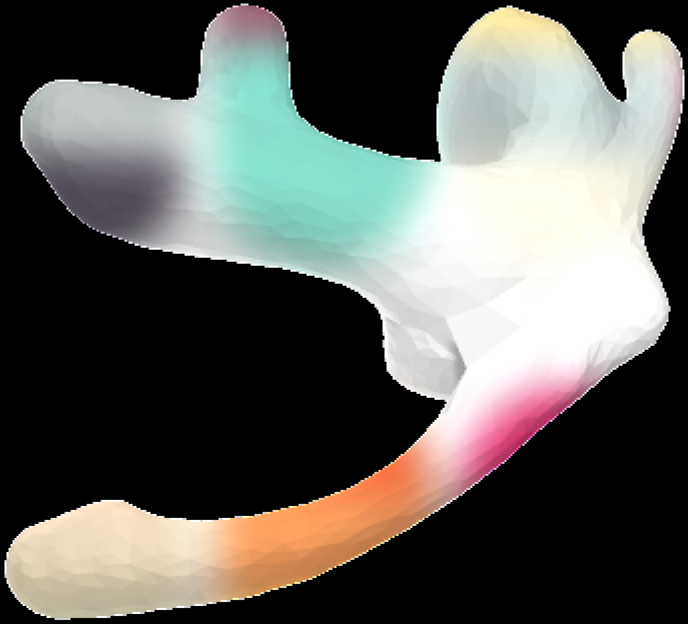

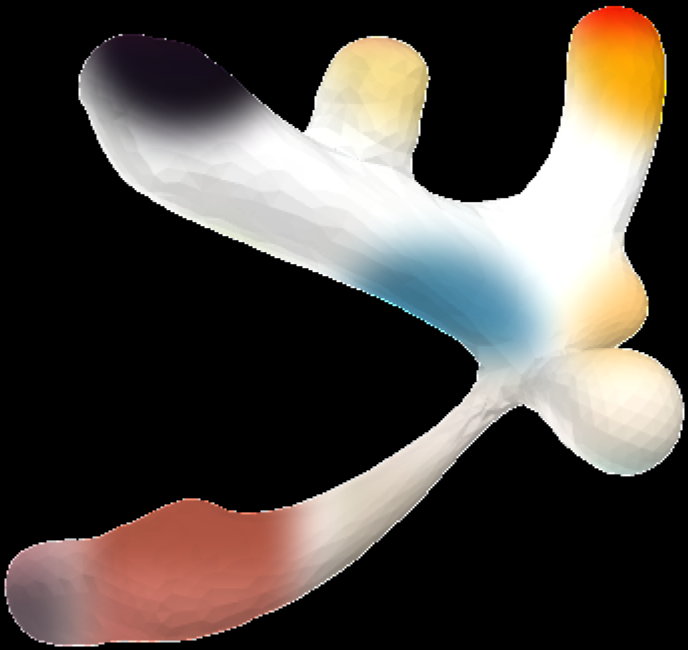

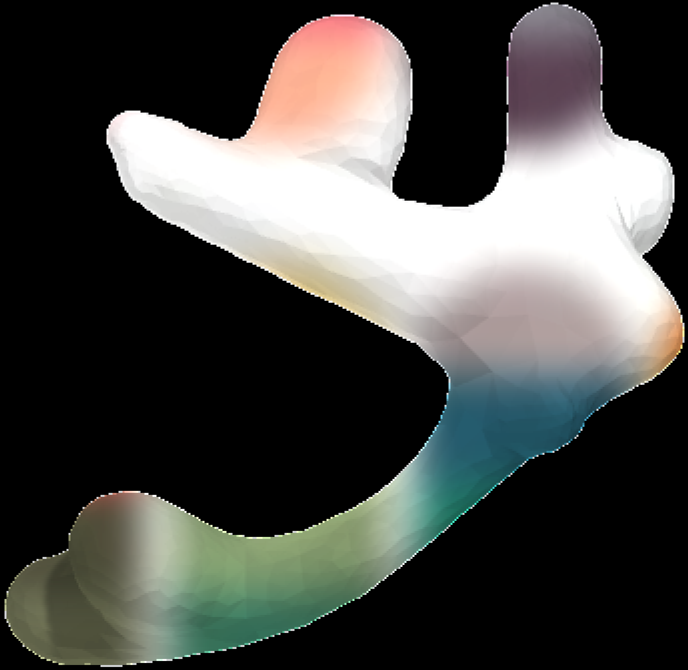

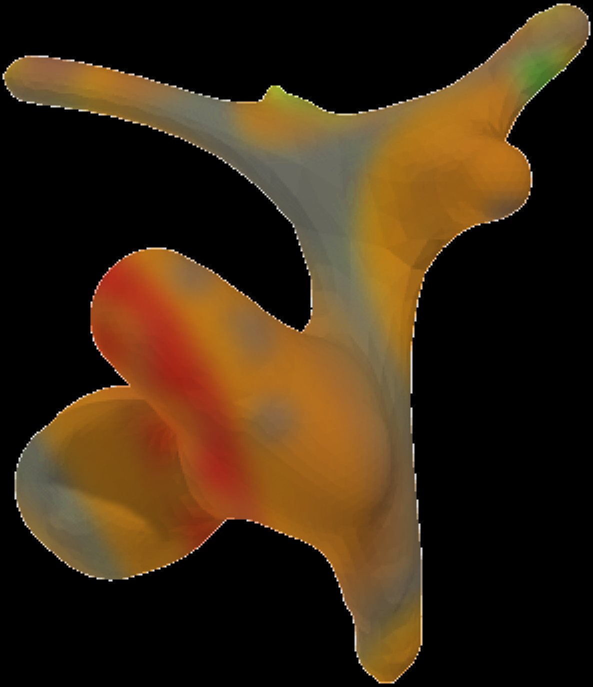

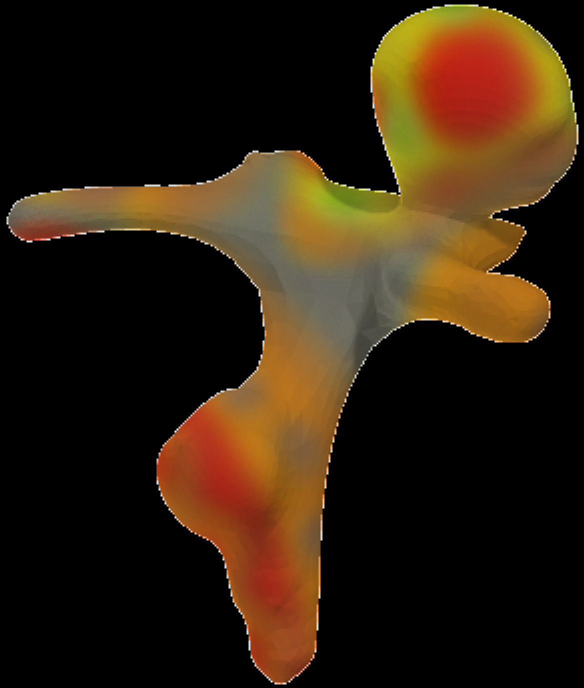

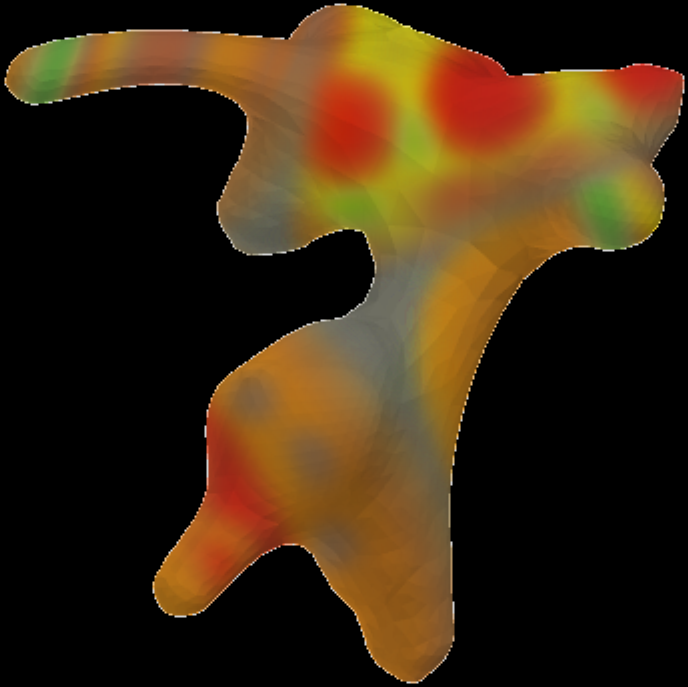

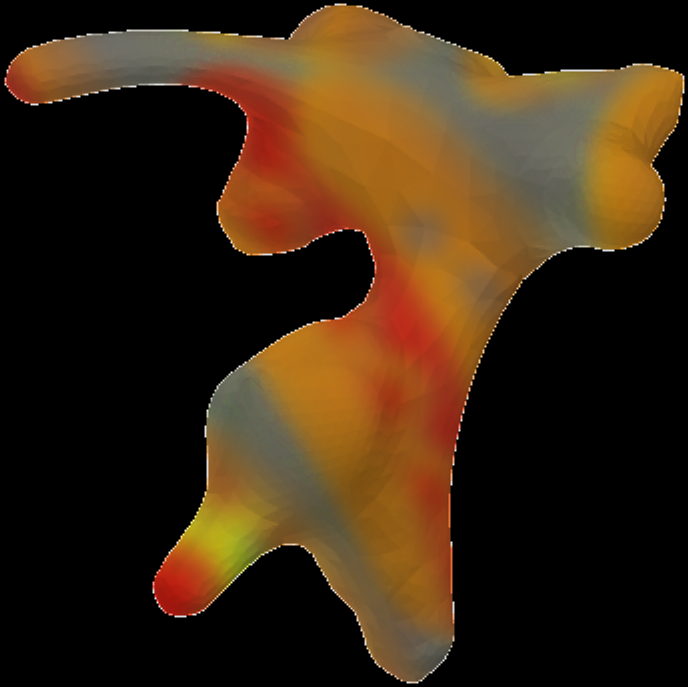

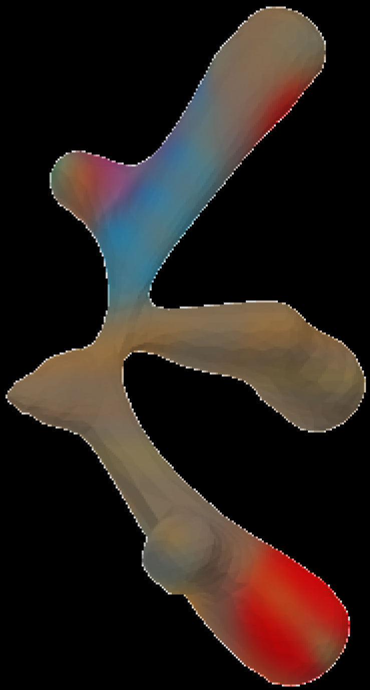

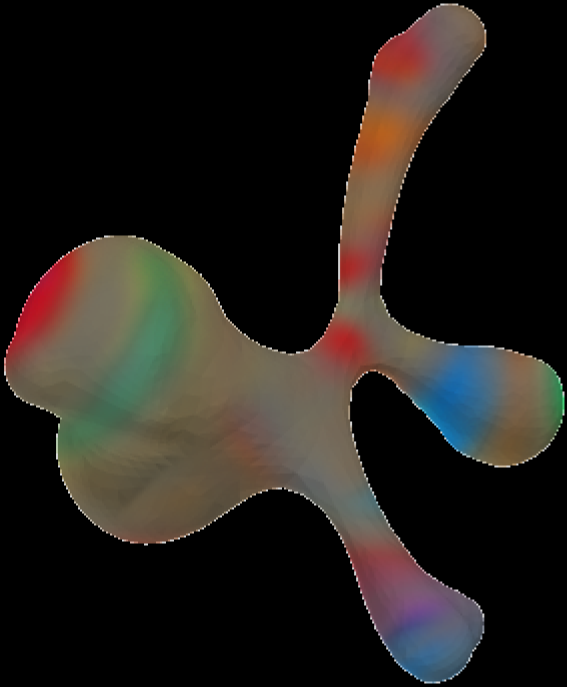

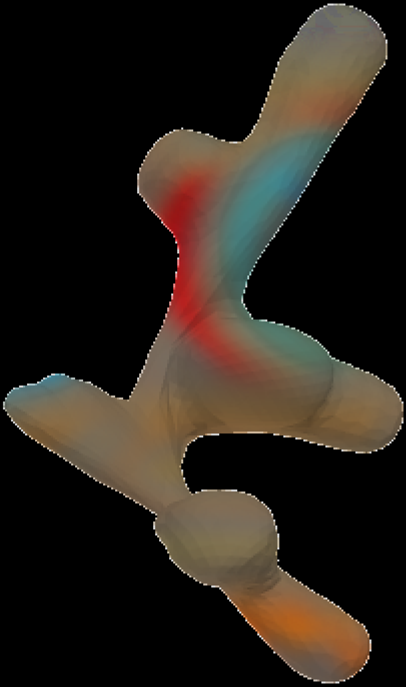

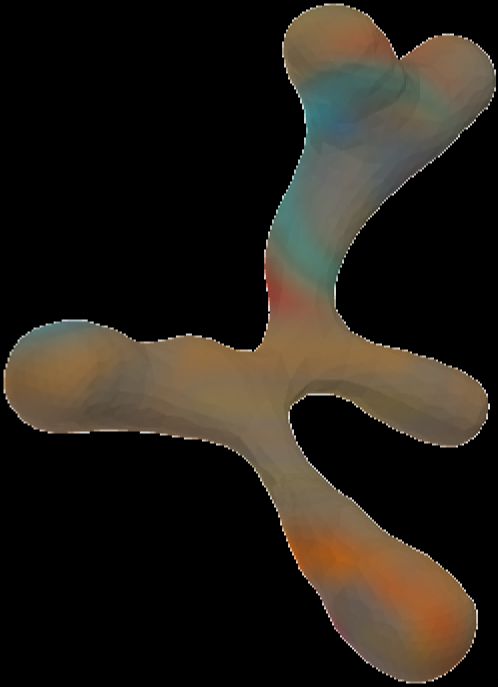

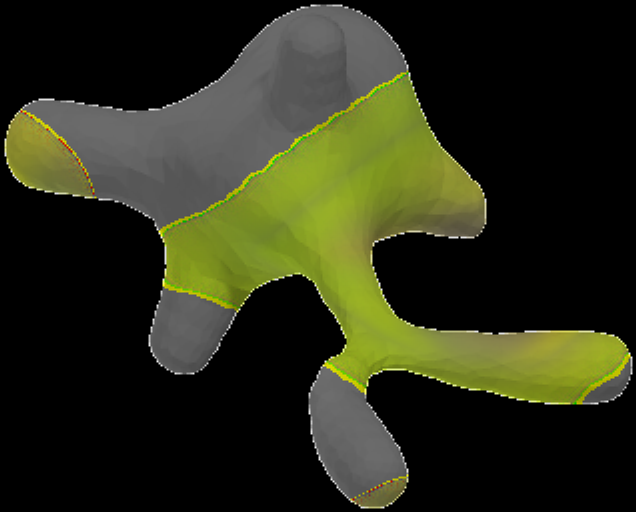

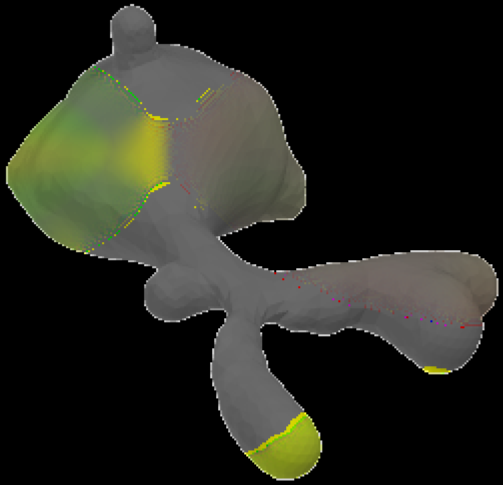

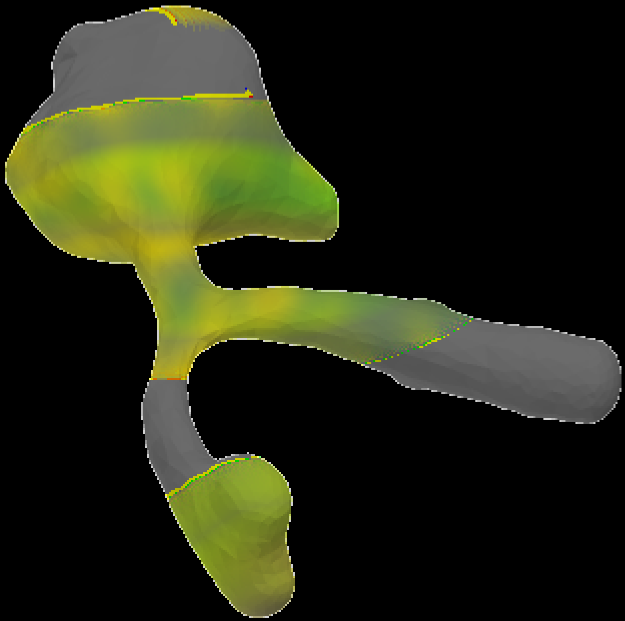

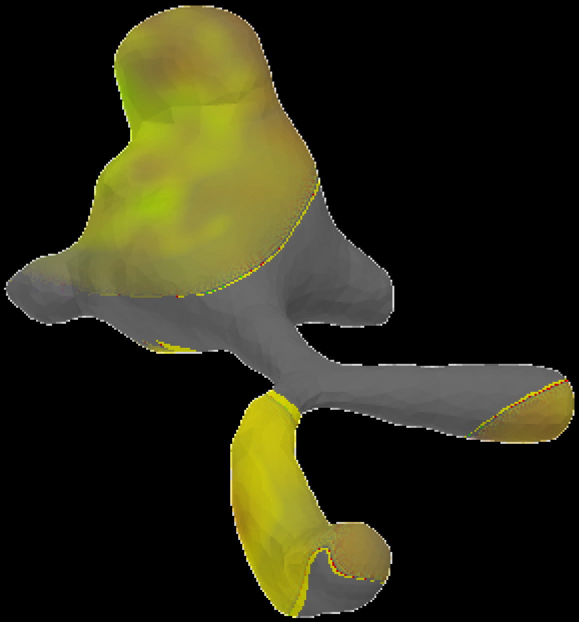

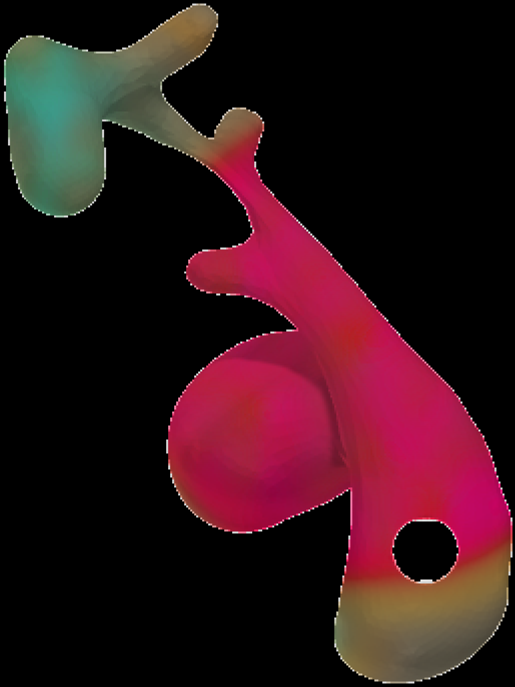

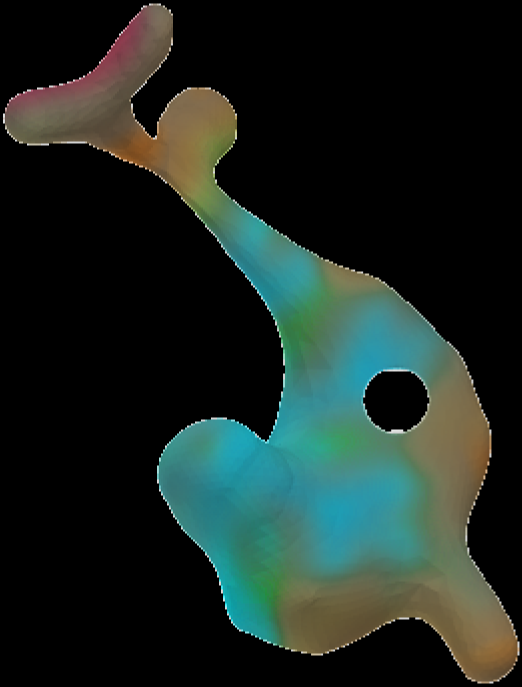

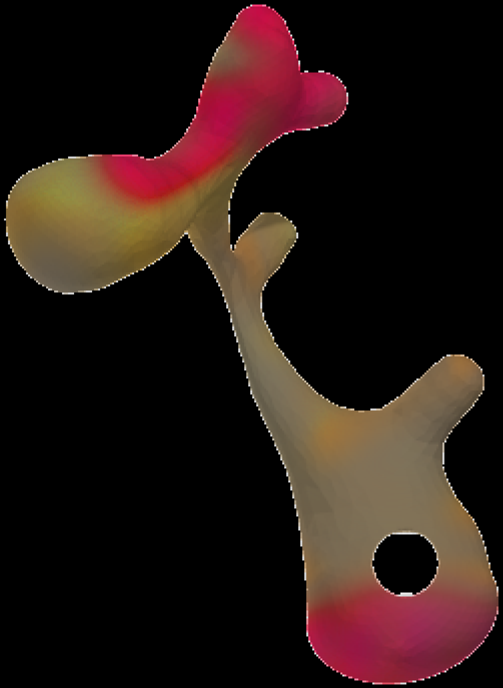

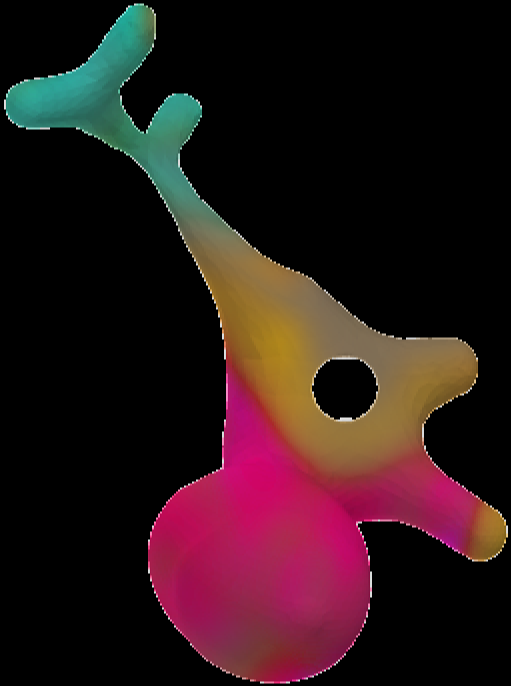

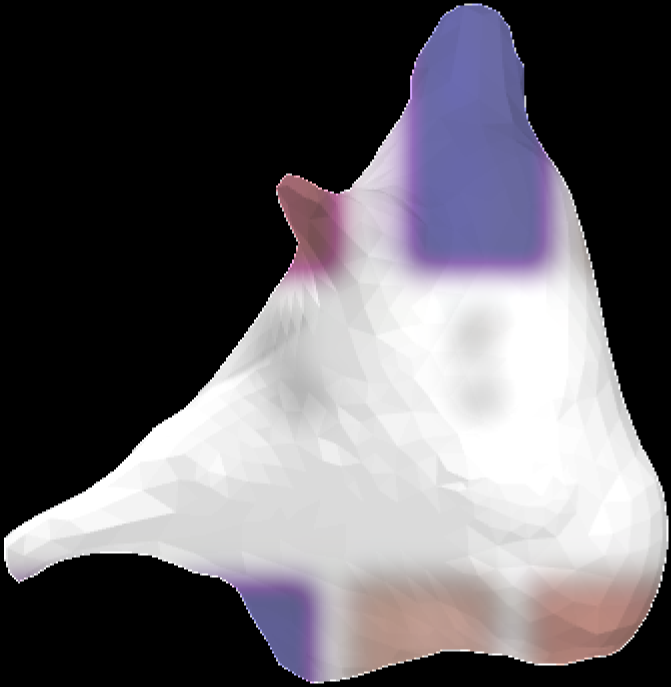

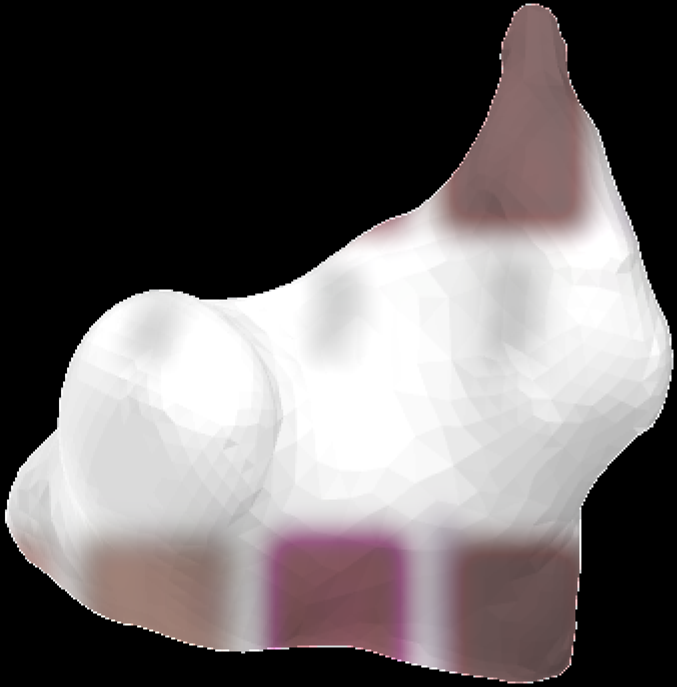

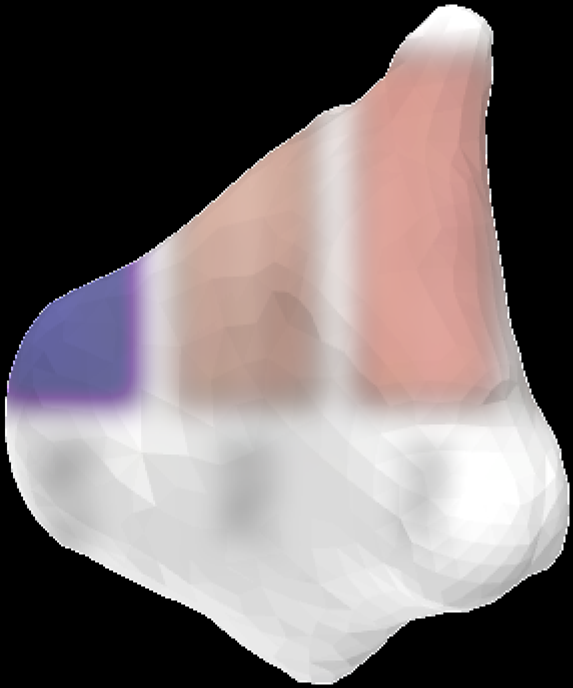

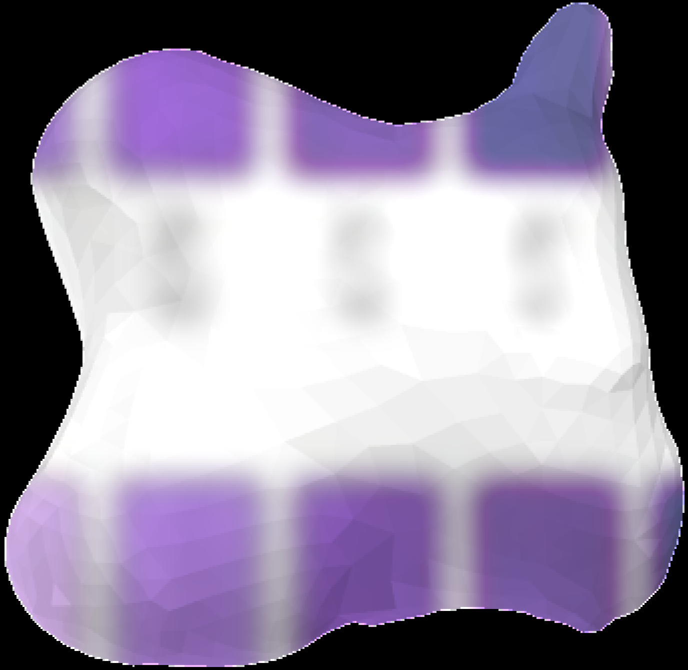

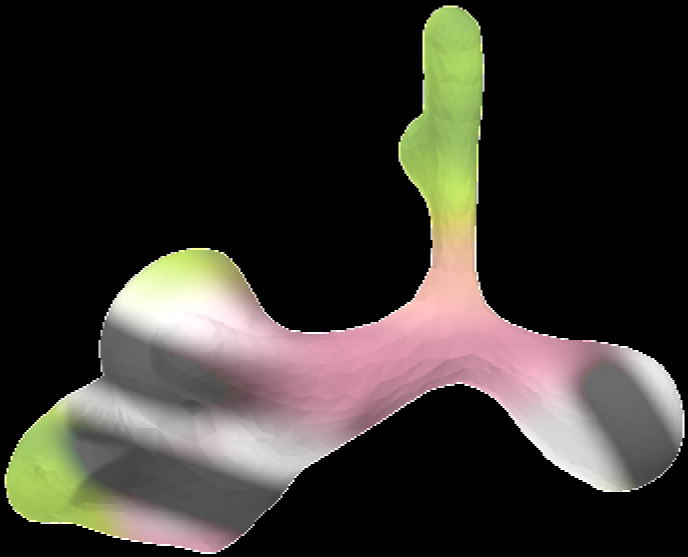

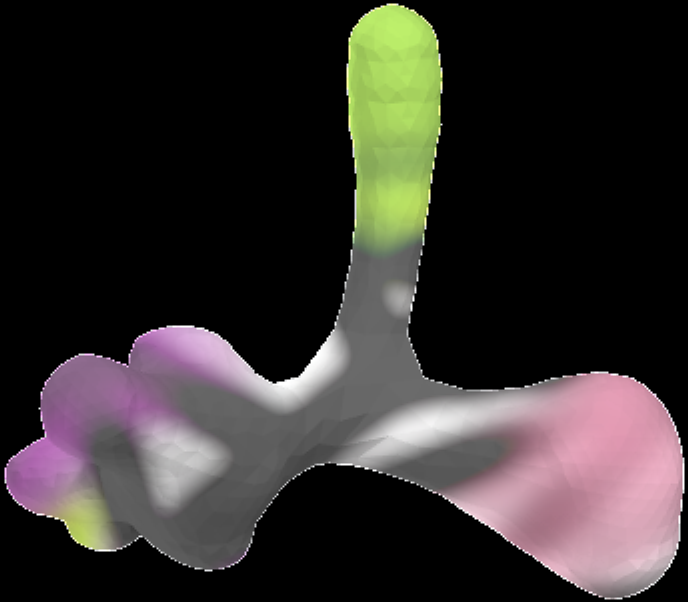

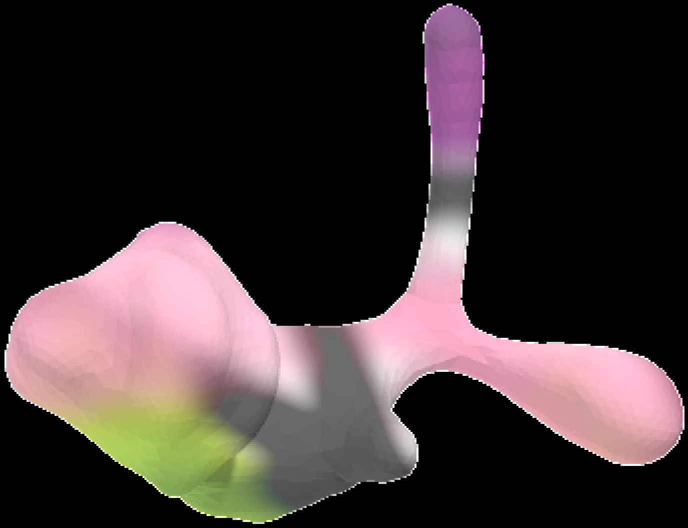

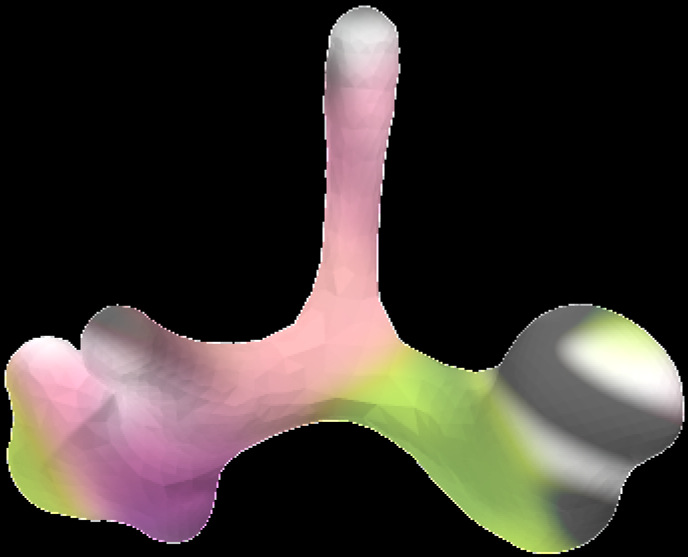

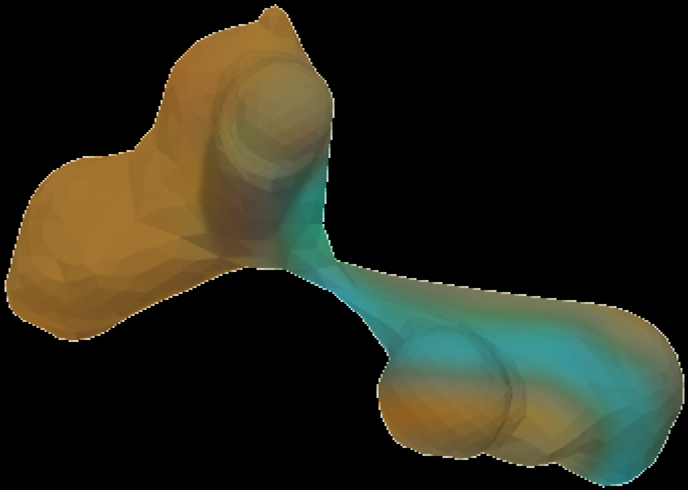

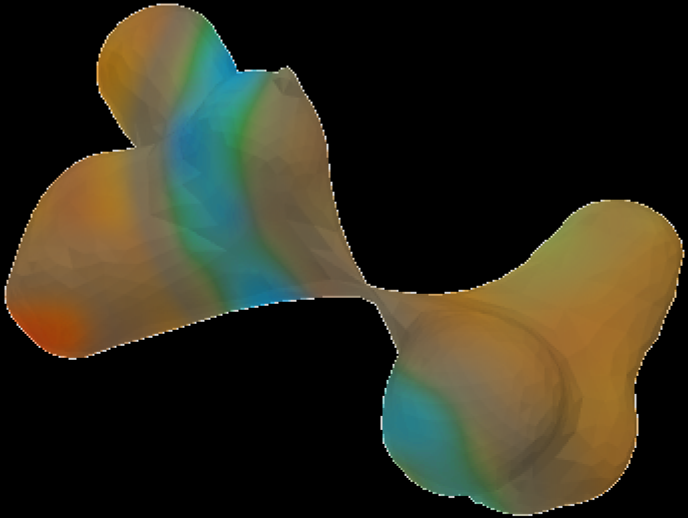

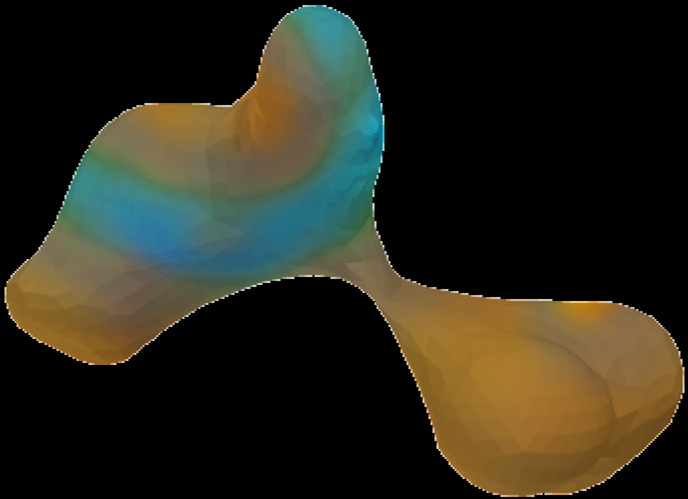

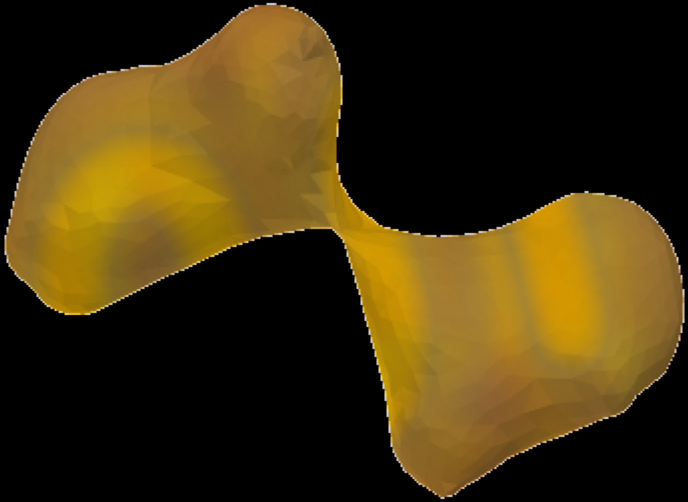

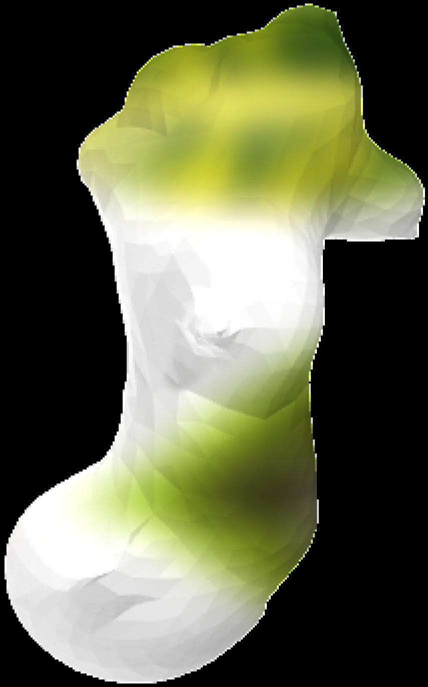

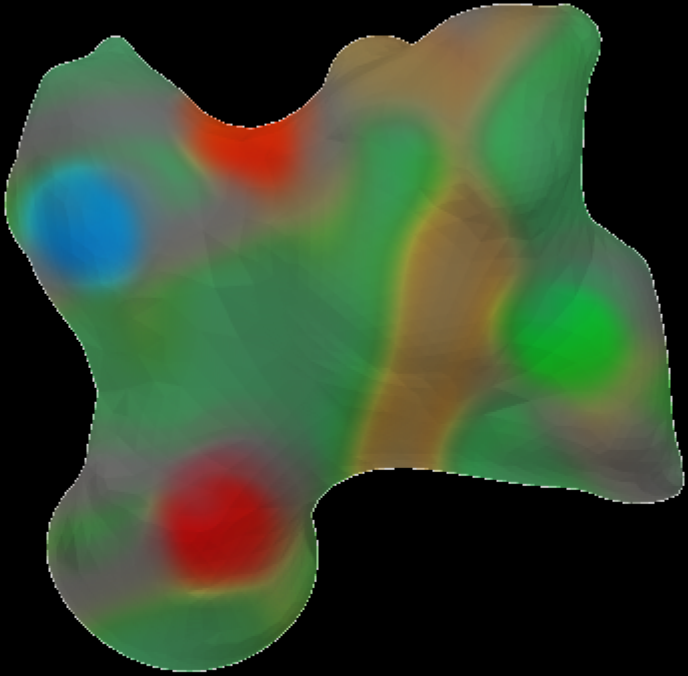

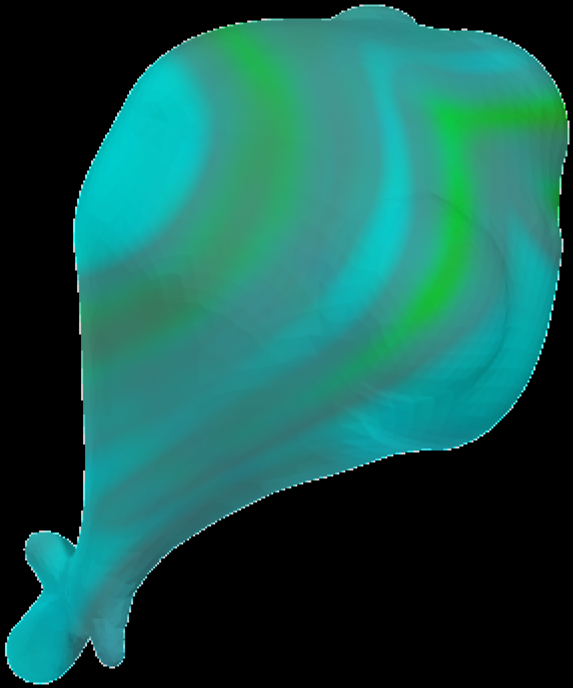

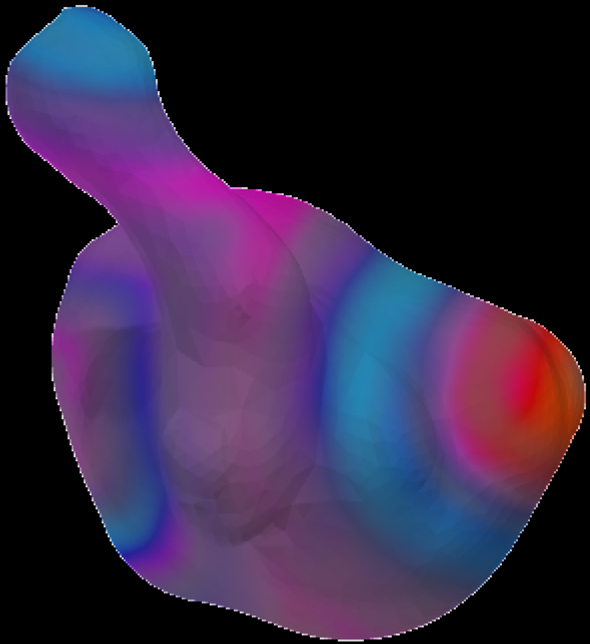

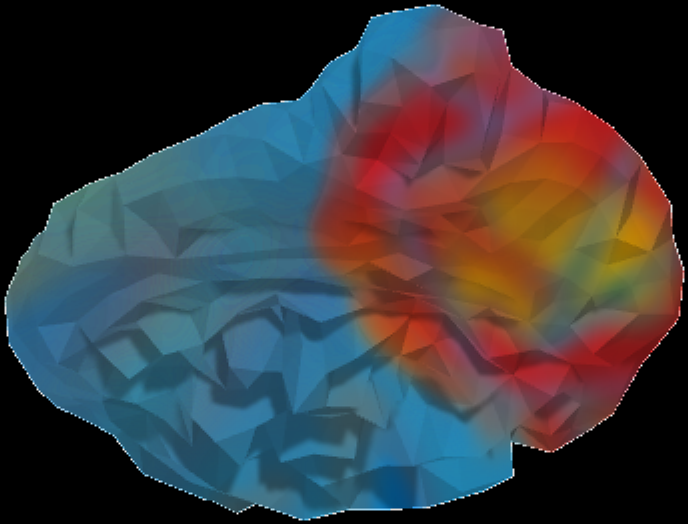

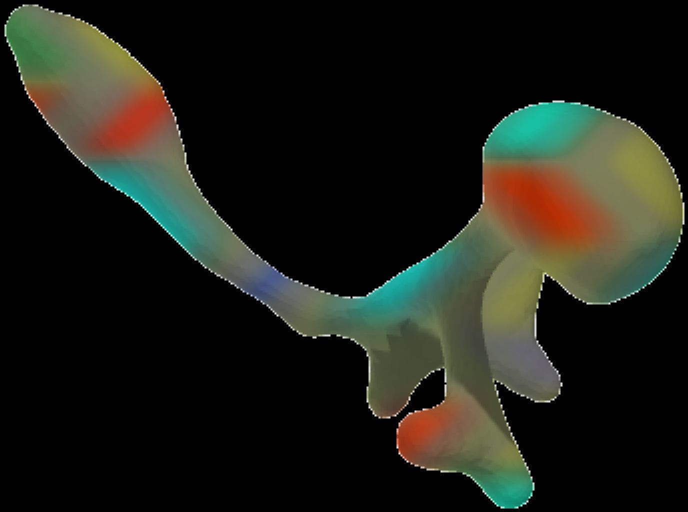

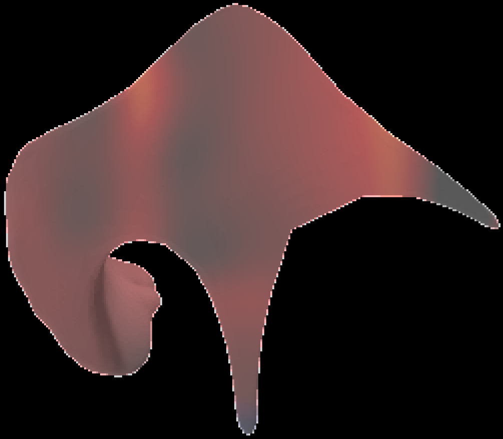

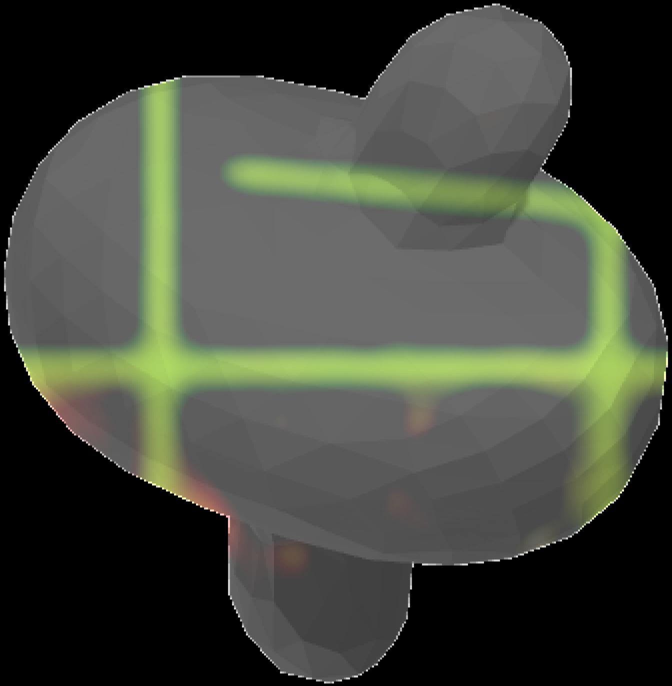

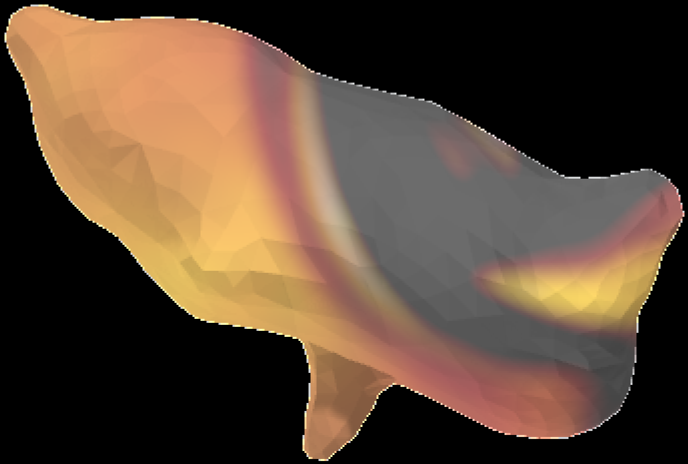

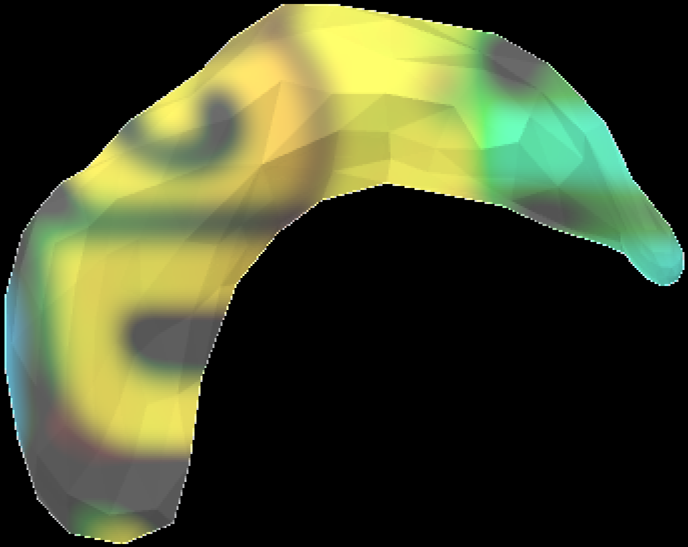

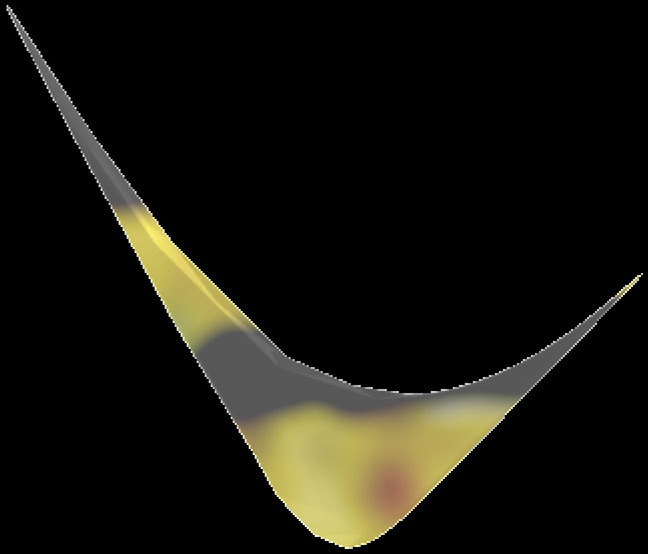

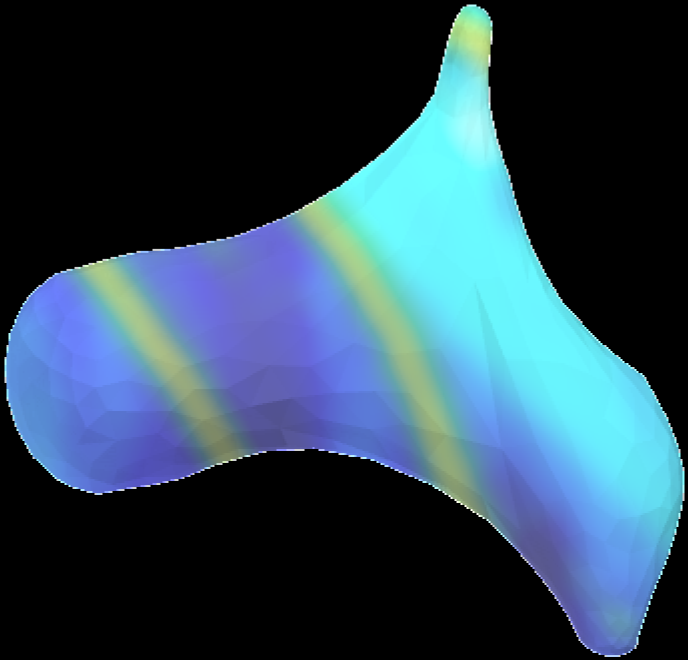

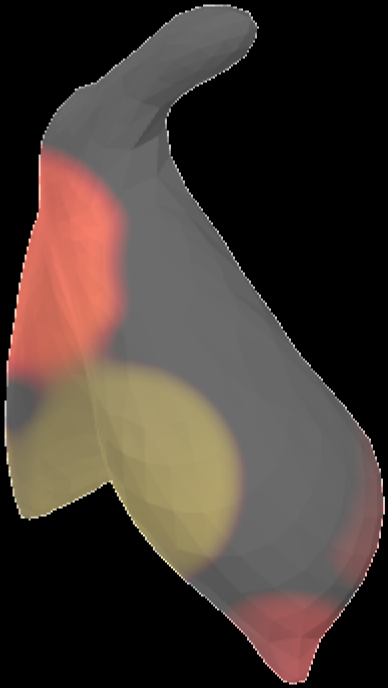

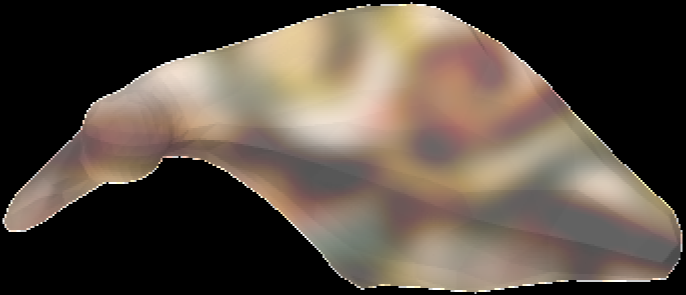

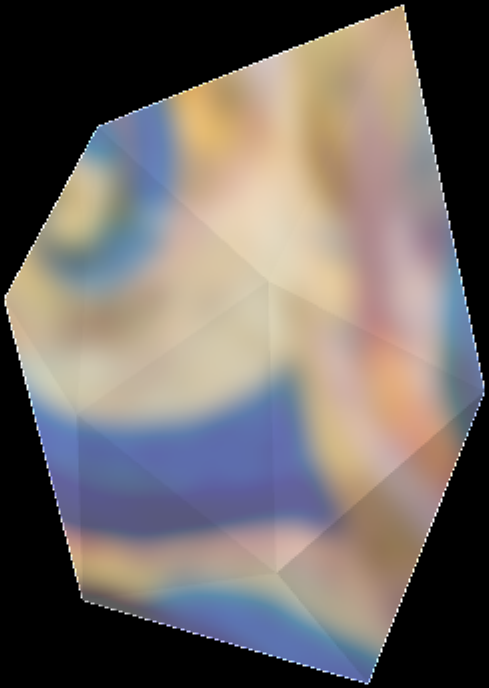

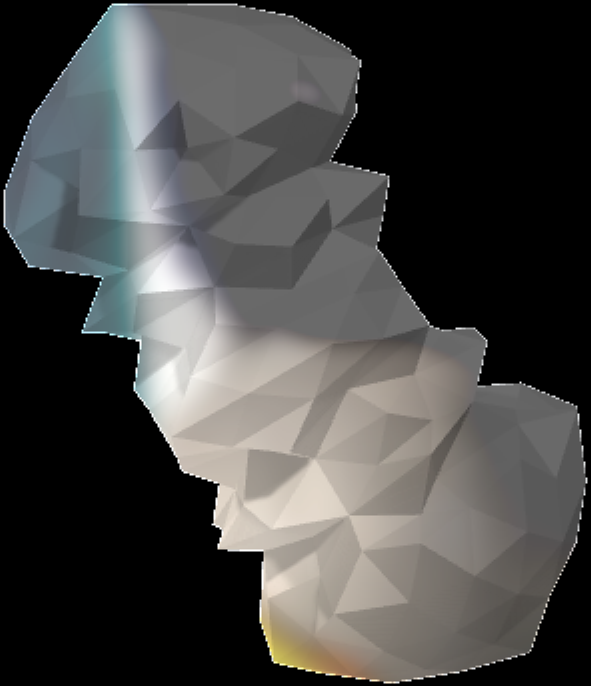

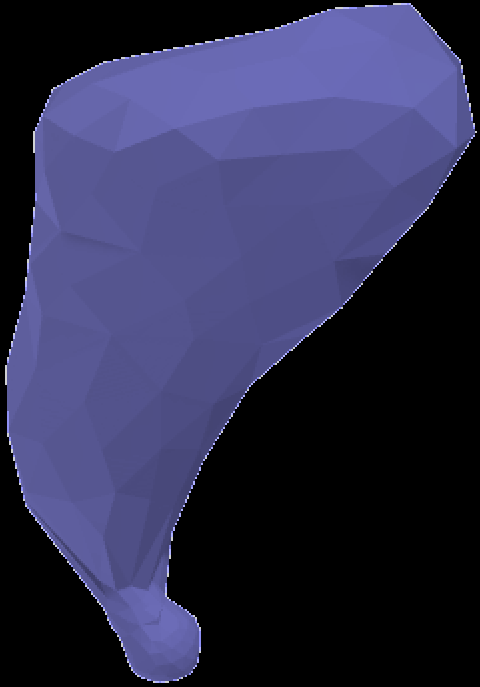

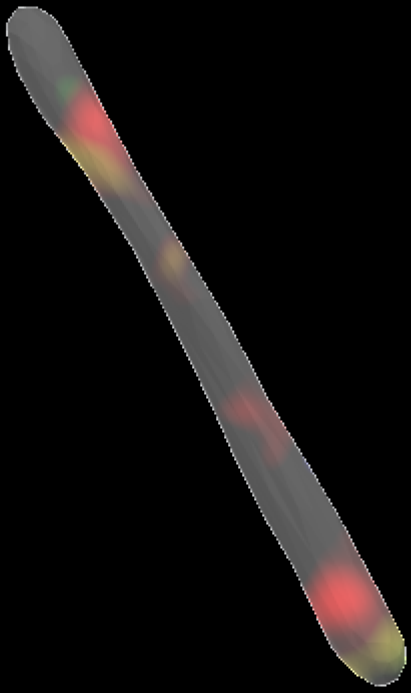

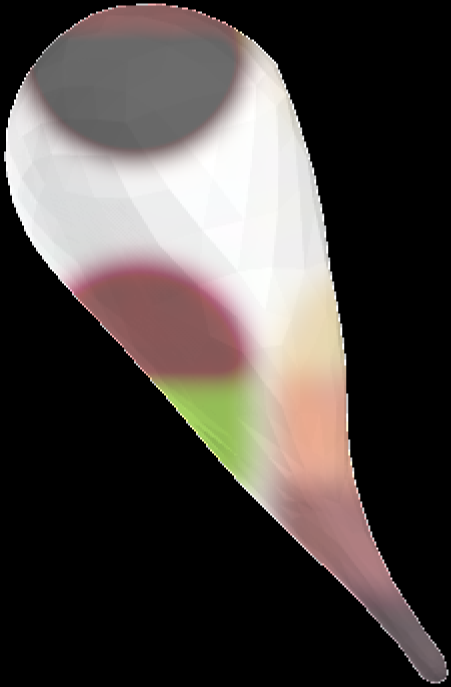

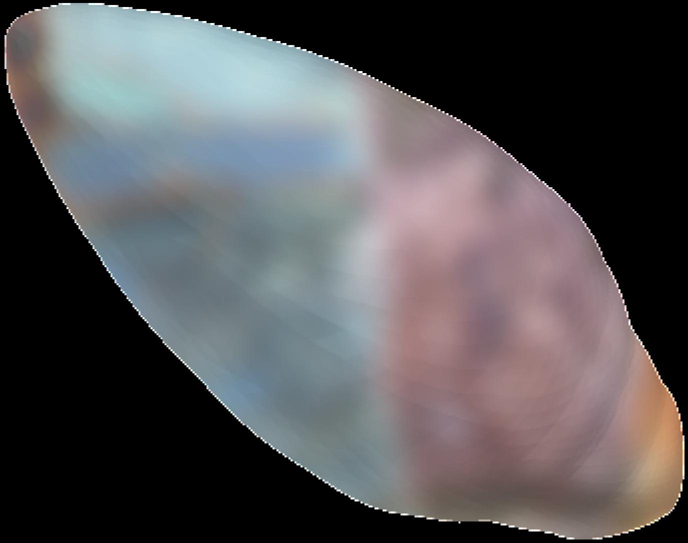

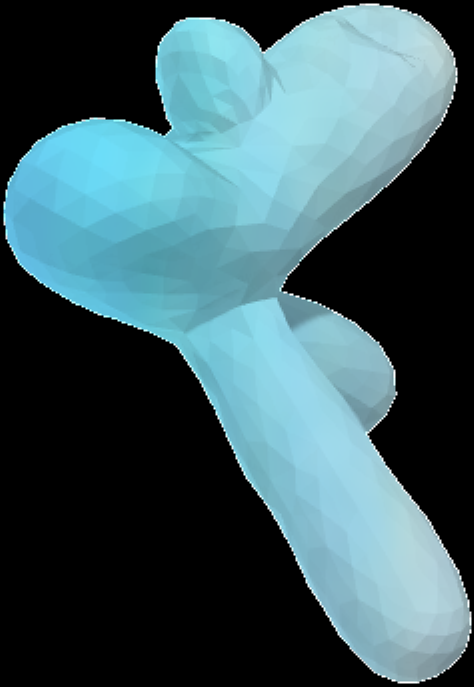

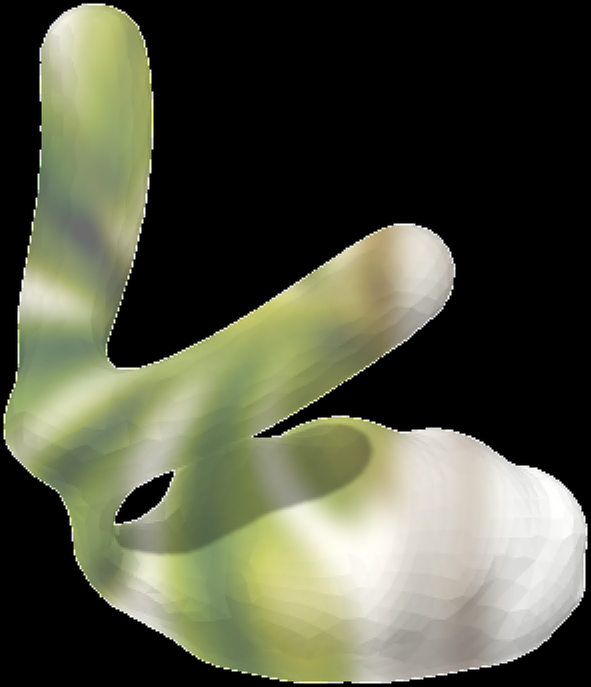

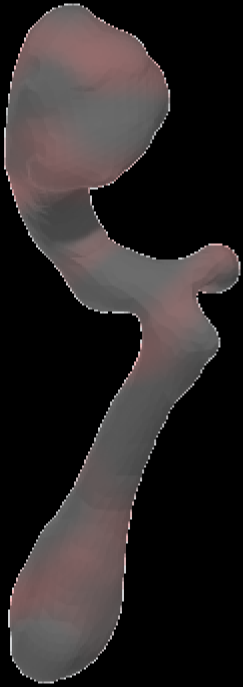

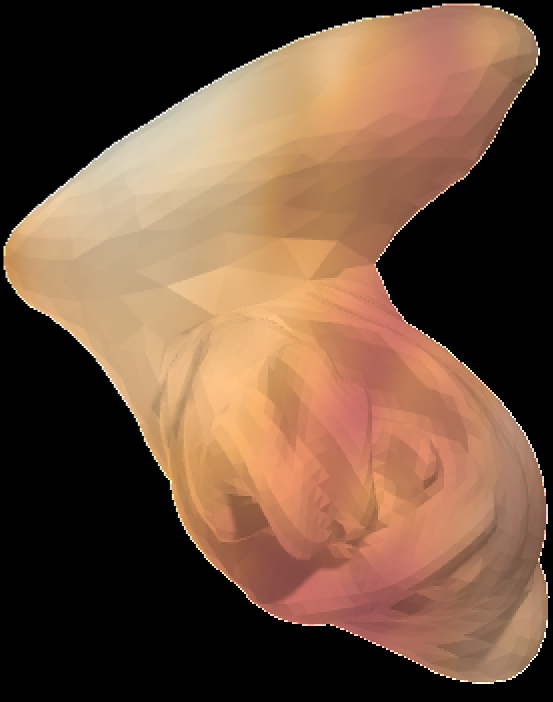

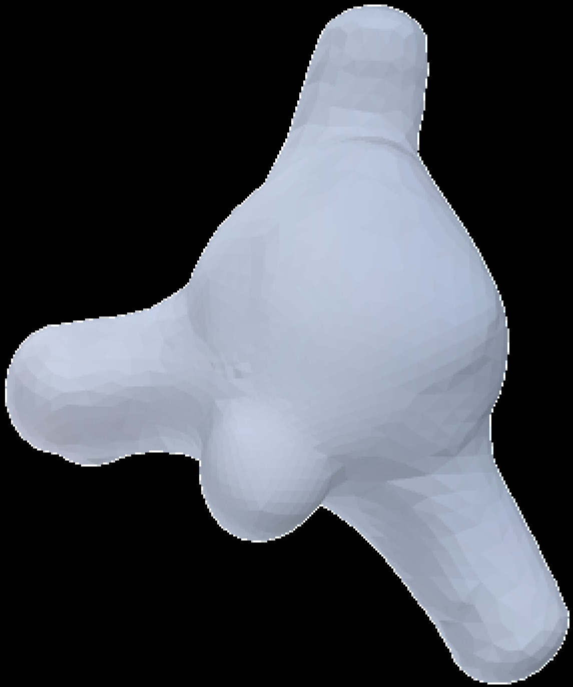

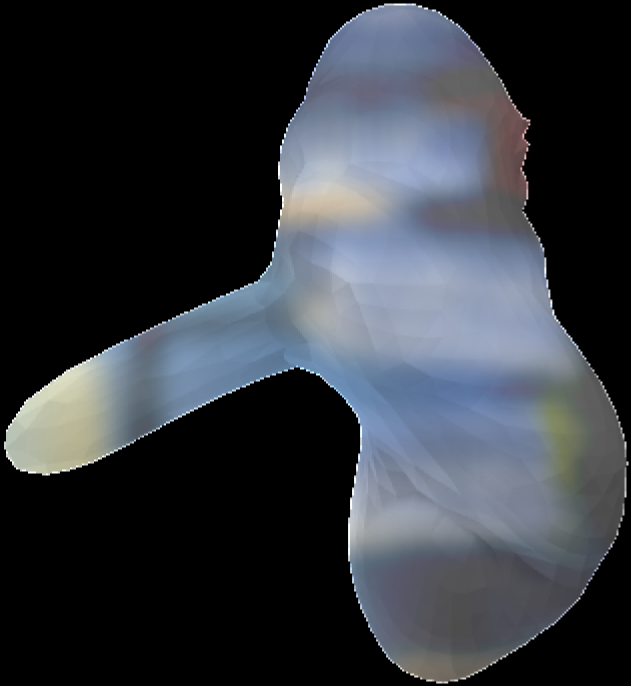

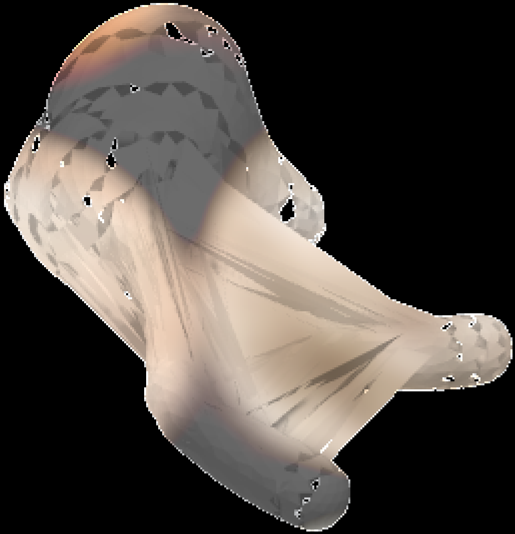

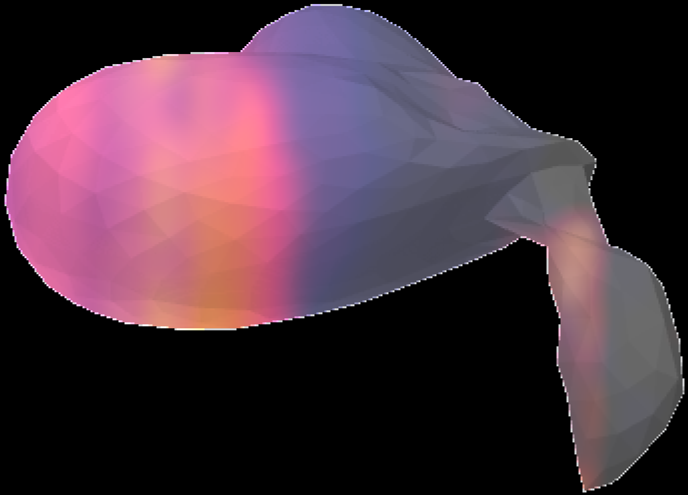

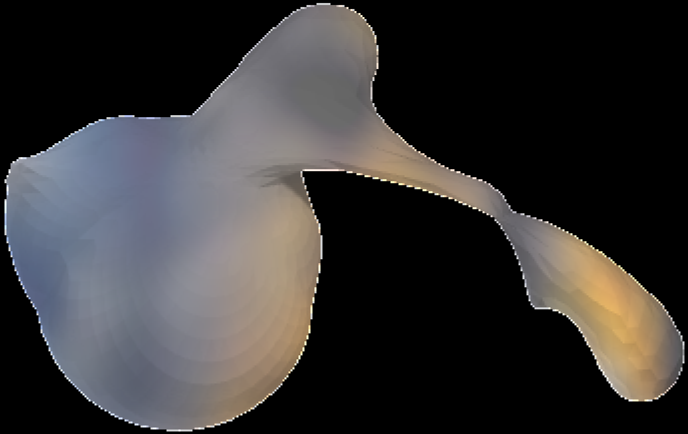

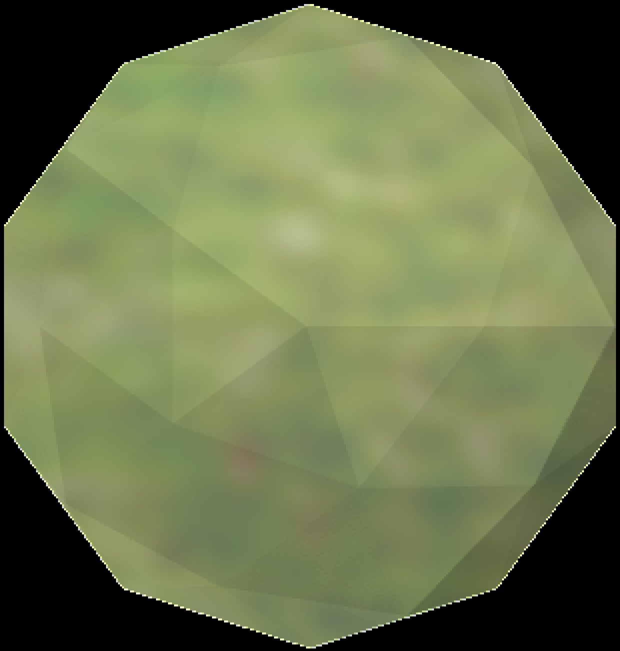

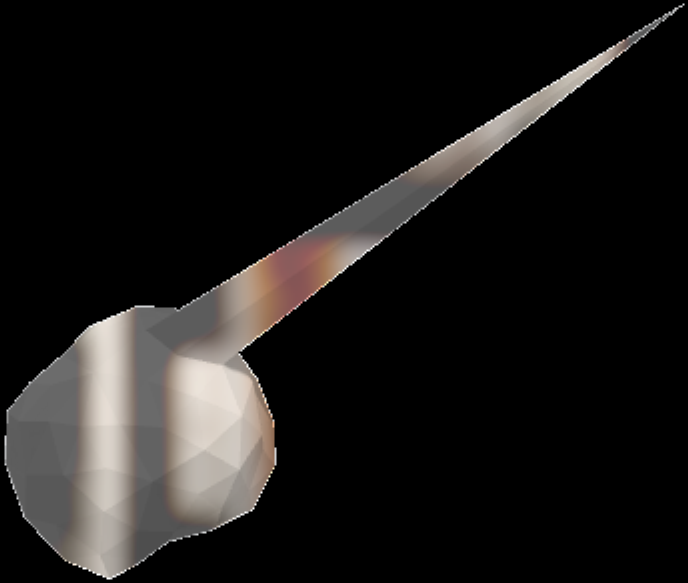
